# Reconstructing an epigenetic landscape using a genetic ‘pulling’ approach

**DOI:** 10.1101/777557

**Authors:** Michael Assaf, Shay Be’er, Elijah Roberts

## Abstract

Cells use genetic switches to shift between alternate stable gene expression states, *e.g.*, to adapt to new environments or to follow a developmental pathway. Conceptually, these stable phenotypes can be considered as attractive states on an epigenetic landscape with phenotypic changes being transitions between states. Measuring these transitions is challenging because they are both very rare in the absence of appropriate signals and very fast. As such, it has proven difficult to experimentally map the epigenetic landscapes that are widely believed to underly developmental networks. Here, we introduce a new nonequilibrium perturbation method to help reconstruct a regulatory network’s epigenetic landscape. We derive the mathematical theory needed and then use the method on simulated data to reconstruct the landscapes. Our results show that with a relatively small number of perturbation experiments it is possible to recover an accurate representation of the true epigenetic landscape. We propose that our theory provides a general method by which epigenetic landscapes can be studied. Finally, our theory suggests that the total perturbation impulse required to induce a switch between metastable states is a fundamental quantity in developmental dynamics.

## Introduction

The presence of many overlapping feedback-based circuits within a cell’s regulatory network has been theorized to give rise to a cellular epigenetic landscape (also called a phenotype landscape) with many metastable states [1, 2]. Fluctuations in the cell’s state due to molecular noise [3–15] randomly drive the cell along this epigenetic landscape bounded by the so-called quasi-potential barriers separating the metastable states. Most of the time the system dwells in the vicinity of one of these metastable states undergoing small random excursions about it. Occasionally, however, a rare, large fluctuation can move the system from one basin of attraction (of a metastable state) to another [16–34].

The stability of these phenotypic states, quantified by the mean first passage time (MFPT) or mean switching time (MST) to transition from one state to another solely via fluctuations, is typically very long to ensure stable phenotypes, and yet cells must transition quickly and deterministically once the proper signal is received [35–38]. Such noise-driven switches, using positive and negative feedback loops, regulate diverse decision-making processes [39–48].

In developmental processes, the regulatory network guides a developing cell through a series of transitions by moving from one metastable state to another along the landscape across the quasi-potential barriers [49–51]. Stochastic fluctuations have been observed to be involved in several developmental processes [52] and developmental transitions may involve quick passage through a number of intermediate states [53]. Indeed, cellular reprogramming under strong perturbations follows a barrier crossing process along a one-dimensional order parameter [54].

Using signals to guide a cell’s state artificially along an epigenetic landscape could open new avenues to treating disease using induced pluripotent stem cells and must also underlie natural differentiation processes [55, 56]. A theory to describe the work that is required to transition a cell between metastable states would be valuable in developing detailed models of differentiation networks as well as designing differentiation protocols. However, reconstructing the cellular epigenetic landscape of a real biological phenotype from steady-state experimental data is usually impossible because of the extreme rareness of the transitions.

Here, we describe a new approach for studying cellular decision landscapes using perturbations. The idea is similar in principle to single-molecule force spectroscopy studies of protein-folding landscapes, allowing one to extract transition information from force-spectroscopy pulling experiments [57, 58]. By pulling a macromolecule or molecular complex at a sufficient force, rare transitions in single molecules such as ligand-receptor dissociation [59], unfolding of a protein [60], or unzipping of nucleic acids [61] can be experimentally observed. The authors in Refs. [57, 58] have devised a theoretical method, in the framework of the Kramers theory, that allows translating the distribution of rupture forces that can be measured experimentally, into the force-dependent lifetime of the system. In our case, starting with a cellular regulatory network, we apply an external “force” or “pulling” to perturb the network in the direction of the desired change. The statistics of the response of the system, *i.e.* the statistics of switching events in the presence of such pulling force, are then used to infer the topology of the landscape, which allows evaluating the lifetime of the various metastable states.

To compute the response of the system to external pulling, we employ a semi-classical approach in the spirit of the Wentzel–Kramers–Brillouin (WKB) theory [29, 33, 62–66] in order to treat the underlying chemical master equation describing the stochastic dynamics of the regulatory network. This formalism allows us to transform the master equation into a set of Hamilton equations which can be dealt with analytically or numerically. We then solve these equations under a prescribed external perturbation with given magnitude and duration and compute the change in the switching probability due to the external pulling. Finally, we use our semiclassical solution in a maximum likelihood framework to infer the parameters of the model from response data. Using the inferred parameters, we reconstruct the epigenetic landscape of the network.

We demonstrate our method on two prototypical model systems: a one-dimensional (1D) system of a self-regulating gene, and a two-dimensional (2D) system of mRNA-protein positive feedback loop. We furthermore discuss how our model can be generalized to higher-dimensional systems.

### Switching in the absence of an external perturbation

Our starting point is an effective 1D model for the dynamics of the protein of interest. It is assumed that the protein is expressed and degraded according to the following set of birth-death reactions

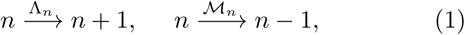

where Λ_*n*_ and *ℳ*_*n*_ are the expression and degradation rates, respectively, and *n* is the current copy number of the proteins.

Neglecting intrinsic noise, the mean number of proteins 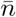 satisfies the following *deterministic* rate equation

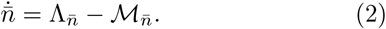

We are interested in a scenario where this rate equation has (at least) three fixed points: *n*_1_ *< n*_2_ *< n*_3_, where *n*_1_ and *n*_3_ are stable fixed points corresponding to the *low* and *high* phenotypes, while *n*_2_ is an intermediate unstable fixed point. One model system that exhibits this property is a protein that positively regulates itself – a self-regulating gene (SRG). While our analysis below is done for generic Λ_*n*_ and *ℳ* _*n*_, in all our simulations we have chosen the birth and death rates to satisfy

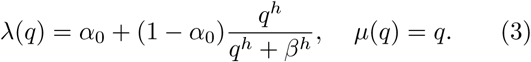

Here *λ*(*q*) = Λ_*n*_*/N* and *µ*(*q*) = *ℳ*_*n*_*/N* are rescaled expression and degradation rates, *q* = *n/N* is the protein density, while *N* is the typical system size, assumed to be large, which represents the typical protein copy number in the *high* state. Furthermore, *α*_0_ is the rescaled base-line expression rate, *h* is the Hill exponent, and *β* is the midpoint of the Hill function. Fig. S1 shows an example of rate equation (2) using rates (3) when the system has three fixed points.

Once intrinsic noise is accounted for, these stable fixed points become metastable, and noise-induced switching between *n*_1_ and *n*_3_ or *vice versa*, occurs. To account for intrinsic noise, we write down the so-called chemical master equation describing the dynamics of *P*_*n*_(*t*) – the probability to find *n* proteins at time *t*:

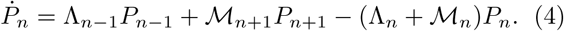

Let us first consider the case of switching in the absence of external perturbations. Here, one can find an exact expression for the mean switching time (MST), by solving the equation for mean first passage time 𝒯 (*n*) –the mean time it takes the system to cross the unstable boundary starting from a state with *n* proteins [67]. Yet, since the analytical solution of this equation is highly cumbersome, and not tractable in the presence of external perturbations, throughout the text we instead use the so-called Wentzel-Kramers-Brillouin (WKB) method [62] to compute the MST, or switching probability.

To set the stage for the WKB method, let us assume without loss of generality that the system starts in the vicinity of the *low* stable fixed point *n*_1_. Assuming the typical system’s size is large, *N* »1, the resulting MST is the MST, expected to be exponentially long, see below. In this case, prior to switching the system enters a long-lived metastable state which is centered about *n*_1_. Indeed, starting from any initial condition *n*_0_ *< n*_2_, after a short 𝒪(1) relaxation time, the dynamics of the probability distribution function can be shown to satisfy the metastability ansatz: *P* (*n ≤ n*_2_, *t*) ≃ *π*(*n*)*e*^−*t/τ*^, while 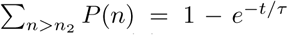 Here *τ* is the MST, *π*(*n*) is called the quasi-stationary distribution (QSD), which determines the *shape* of the metastable state, and it is evident that the probability to be at *n > n*_2_ is negligibly small at times *t* ≪ *τ*.

We now plug this ansatz into master equation (4), and neglect the exponentially small term proportional to *τ* ^−1^ (see below). Employing the WKB ansatz *π*(*n*) ≡ *π*(*q*) ∼ exp[*NS*(*q*)] on the resulting quasistationary master equation, where *S*(*q*) is the action function, yields a stationary Hamilton-Jacobi equation *H*(*q, ∂*_*q*_*S*) = 0, with the Hamiltonian being

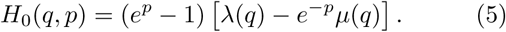

Here *p* = *∂*_*q*_*S* is called the momentum in analogy to classical mechanics, while the subscript 0 stands for the unperturbed case. To find the optimal path to switch – the path the system takes with an overwhelmingly large probability during a switching event [70, 71] – we need to find a nontrivial heteroclinic trajectory, *p*_0_(*q*), connecting the saddles (*q, p*) = (*q*_1_, 0) and (*q*_2_, 0) [62–66]. Equating *H*_0_ = 0 yields

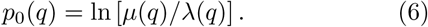

Thus, the action function is found by integrating: *S*(*q*) = *p*(*q*^*′*^)*dq*^*′*^. Given the initial metastable state *q*_1_ = *n*_1_*/N* and unstable fixed point *q*_2_ = *n*_2_*/N*, the MST between the *low* and *high* states can be shown to satisfy in the leading order [62, 64, 65, 69]

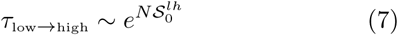

where 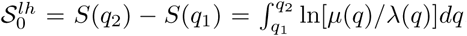, is the switching barrier between the *low* and *high* states, in the absence of an external force. Similarly, 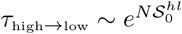, where 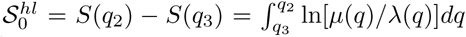, is the switching barrier between the *high* and *low* states. For *N* ≫ 1, these MSTs are indeed exponentially large thus validating our a-priori metastability assumption (see Fig. S2). Note, that in the absence of external perturbation, the pre-factor of *τ* can be accurately found as well [65, 72].

In the following, rather than the MST, we will be interested in computing *𝒫*^*lh*^ and *𝒫*^*hl*^ – the switching *probabilities* over some time *t* «*τ* starting from the *low* to *high* and *high* to *low* states, respectively. For example, starting from the vicinity of *n*_1_, *𝒫*^*lh*^ is determined by the fraction of stochastic realizations of process (1) that cross *n*_2_ in a given time *t* out of the total number of realizations. Using the metastability ansatz, and demanding that the total probability be unity, we have 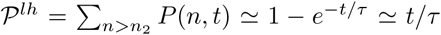, where the last approximation holds for *t* ≪ *τ*; that is, *𝒫*^*lh*^ is exponentially small at *t* ≪ *τ*. As a result, in the absence of external force, and using a similar argument for the calculation of *𝒫*^*hl*^, the switching probabilities up to some arbitrary time *t* ≪ *τ* satisfy in the leading order

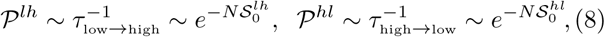

where logarithmic corrections depending on the arbitrary time *t* and the pre-factor entering *τ* have been omitted.

### Switching in the presence of an external perturbation

#### Low-to-high switch

Let us begin by studying the case of *low* to *high* switch in the presence of an external perturbation. To do so, we add an external time-dependent force to the protein’s expression rate, Λ_*n*_ → Λ_*n*_ + *ϕ*(*t*), where *ϕ*(*t*) is applied for a finite duration *T* such that

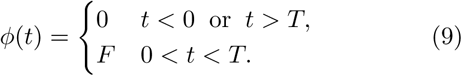

As can be seen in Fig. S3, the result of this perturbation is that the system is pushed nearer to the switching barrier and with some increased probability can then switch to the *high* state. The switching probability depends on both the force *F* and the duration *T* of the perturbation.

Note that, in general, the system does not need to relax to a new quasi-stationary distribution during the perturbation. We are interested in computing the dependence of the change in the *low* to *high* switching probability *𝒮*^*lh*^ on both *F* and *T*.

Given the time-dependent protocol *ϕ*(*t*) [Eq. (9)] for the change in the protein’s expression rate, one can perform a similar WKB analysis as done above in the unperturbed case. This yields two distinct Hamiltonians: the unperturbed Hamiltonian (5) before and after the external perturbation has been applied, and the Hamiltonian during the perturbation with an elevated expression rate:

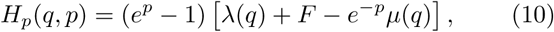

where the superscript *p* stands for the perturbed case. Each of the two Hamiltonians is an integral of motion on the corresponding time interval. Here, the optimal switching path [*q*_*op*_(*t*), *q*_*op*_(*t*)] starts at the saddle point (*q, p*) = (*q*_1_, 0) well before the perturbation has been applied, and ends at the saddle point (*q, p*) = (*q*_2_, 0), well after the perturbation has been applied [73, 74]. It can be found by matching three separate trajectory segments: the pre-perturbation, perturbation, and post-perturbation segments (see Fig. 1a).

**Figure 1.**
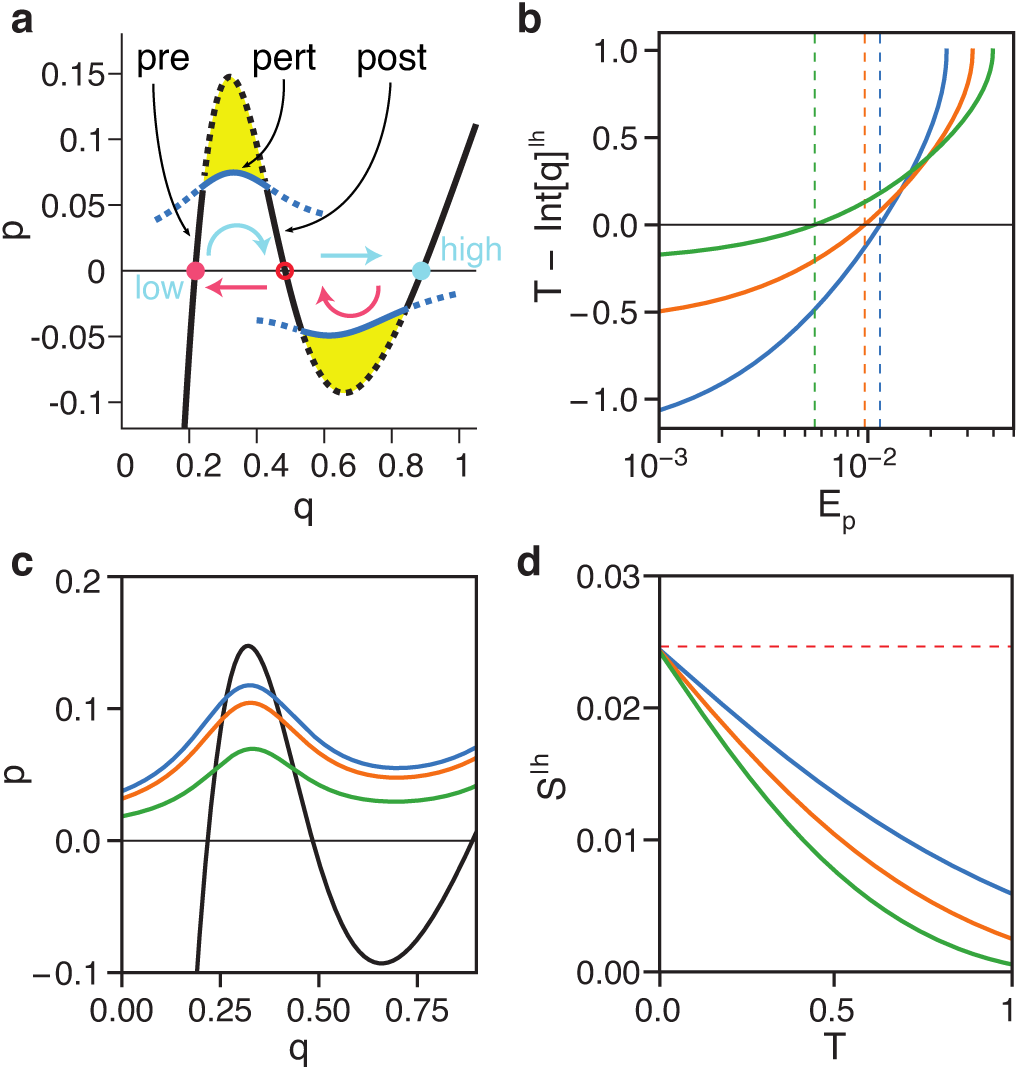
Calculation of the optimal switching path for the self-regulating gene. (a) Illustration of the optimal paths to switching without (black) and with (blue) perturbation. The path segments are labeled as pre-perturbation (pre), perturbation (pert), and post-perturbation (post), see text. The top path shows *low* to *high* switching and the bottom path shows *high* to *low*. The yellow areas give the decrease in the momentum barriers due to the perturbation. In the *low* to *high* switch we have taken *F* = 0.15 and *E*_*p*_ = 0.01 such that *T* = 1.1, whereas for the *high* to *low* we have taken *F* = 0.2 and *E*_*p*_ = 0.005 such that *T* = 2.06. The other parameters are *α*_0_ = 0.2, *β* = 0.562 and *h* = 4. (b) Plot of 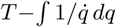 [see equation (12)] vs *E*_*p*_ for the *low* to *high* switch under three different values of the perturbation strength *F* : 0.15 (blue), 0.20 (orange), 0.25 (green). Also shown are (c) momentum *p* vs coordinate *q* and (d) the change in action Δ*S* vs perturbation time *T* for the same three *F* values.

The matching conditions at times *t* = 0 and *t* = *T* are provided by the continuity of the functions *q*(*t*) and *p*(*t*) [73, 74]. The pre- and post-perturbation segments must have a zero energy, *E* = 0, so they are parts of the original zero-energy trajectory, *p*_0_(*q*), see Eq. (6). Yet, for the perturbation segment, the energy *E* = *E*_*p*_ is nonzero and a-priori unknown. It parameterizes the intersection points 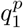 and 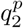 between the unperturbed zero-energy line *p*_0_(*q*) [Eq. (6)] and the perturbed path, *p*_*p*_(*q*) [73, 74]. The latter is the solution to the equation *H*_*p*_(*q, p*) = *E*_*p*_, which is given by:

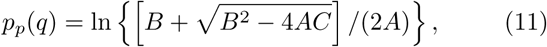

where *A, B* and *C* are functions of *q* and satisfy *A* = *λ*(*q*) + *F*, *B* = *λ*(*q*) + *F* + *µ*(*q*) + *E*_*p*_, and *C* = *µ*(*q*).

To determine the energy *E*_*p*_, we demand that the duration of the perturbation be *T* [73, 74]. Thus, we have:

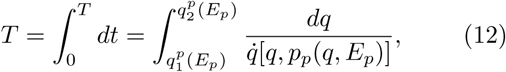

where 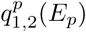 are the intersection points between the unperturbed *p*_0_(*q*) and perturbed *p*_*p*_(*q*) trajectories, and 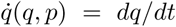 is given by Hamilton’s equation 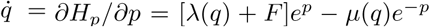. Therefore, plugging *p*_*p*_(*q*) from Eq. (11) into 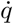, Eq. (12) becomes:

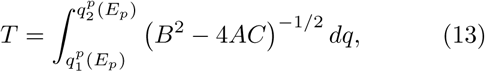

where *A, B* and *C* are given below Eq. (11). Putting it all together, using the fact that the action satisfies 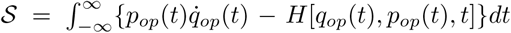 [75, 76], and recalling that *dS* = (*∂S/∂t*)*dt* + (*∂S/∂q*)*dq*, we finally arrive at the corrected switching barrier from the *low* to *high* states:

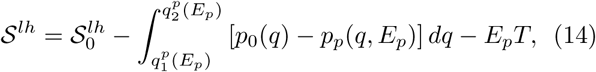

where *E*_*p*_ = *E*_*p*_(*F, T*) can be found from Eq. (13), 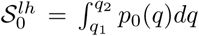 is the unperturbed switching barrier from the *low* to *high* states, and we have used the fact that 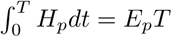.

Note that, for the birth and death rates of the SRG model (3), Eq. (13) has no closed form solution. To study the switching behavior under such perturbation, we first numerically evaluate the integral equation to find the matching *E*_*p*_ (see Fig. 1b). Given a numerical value for *E*_*p*_, we then use Eq. (14) to calculate the perturbed action (Fig. 1c-d). Finally, we use Eq. (8) to calculate the perturbed switching probability *𝒫* ^*lh*^.

To evaluate the accuracy of our theory, we compared the dependence of *𝒫*^*lh*^ on *F* and *T* between theory and Monte Carlo simulations [77]. We calculated *𝒫*^*lh*^ for a range of *F* values for three different perturbation times *T*. Because we are trying to develop a theory that is directly relatable to biological experiments, we limited the range of *F* values to those with *P* ^*lh*^ *>* 1 ×10^−6^. Detecting a cell phenotype with a frequency of one per million cells is at the limit of feasibility using flow cytometry techniques. We also limited the comparison to *𝒫* ^*lh*^ *<* 1 ×10^−2^, below which the switching barrier *𝒮* ^*lh*^ starts to become low enough such that the WKB approximation is invalidated [65]. As can be seen in Fig. 2a, there is excellent agreement between theory and numerics.

**Figure 2.**
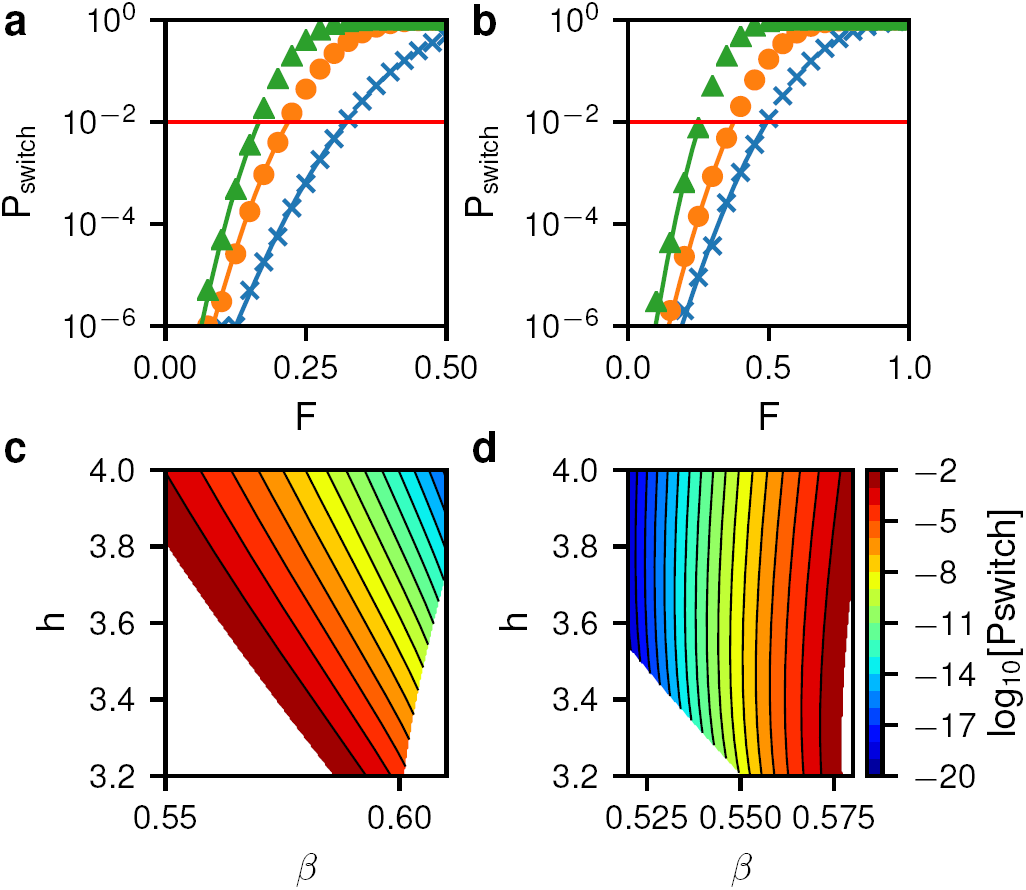
Change in the switching probability with perturbation. (a) Change in switching probability vs perturbation strength *F* for the *low* to *high* switch. Symbols and lines give numerical and theoretical values, respectively. A constant prefactor of 0.15 was used with the theory. Data are shown for three values of the perturbation time *T* : 0.5 (blue ×), 0.75 (orange ◦), 1.0 (green Δ). (b) Change in switching probability vs perturbation strength for the *high* to *low* switch with perturbation times 0.75 (blue ×), 1.0 (orange ◦), and 1.5 (green Δ), and a prefactor of 0.2. (c+d) Switching probabilities in the *β* vs *h* plane for the *low* to *high* switch (c) and the *high* to *low* switch (d).

Finally, we note that our result for the increase in the switching probability in the aftermath of an external perturbation [Eq. (14)] can be simplified in three particular limits: (i) close to the bifurcation limit, where either the *low* and intermediate, or the *high* and intermediate fixed points merge and the switching barrier vanishes, (ii) for weak external force, *F*≪ 1, and (iii) in the case of *h* → *∞*, *i.e*, a very steep regulatory function. Close to the bifurcation limit, we find that *𝒮* ^*lh*^ depends only on the *impulse* of the perturbation, *FT*, see Discussion and Appendix A; in the case of weak force, we show that the increase in the switching probability is exponential in *F*, see Appendix B, while in the limit of *h* → *∞*, where the expression rate is given by a heaviside step function, we find an explicit expression for *𝒮* ^*lh*^ as function of *F* and *T*, see Appendix C.

#### High-to-low switch

We now briefly describe the case of switching from the *high* to *low* states in the presence of an external perturbation. Here, switching can be driven by increasing the protein’s degradation rate. That is, we have *µ*(*n*) →*µ*(*n*)[1 + *ϕ*(*t*)], where *ϕ*(*t*) is given by Eq. (9).

Following along the same lines as the previous subsection, here the perturbed Hamiltonian becomes

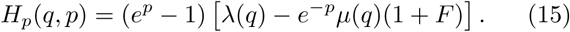

As a result, the intersection points 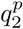 and 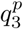 are now determined by equating *p*_0_(*q*) from Eq. (6) with the perturbed path *p*_*p*_(*q*), given by Eq. (11), with *A* = *λ*(*q*), *B* = *λ*(*q*) + *µ*(*q*)(1 + *F*) + *E*_p_, and *C* = *µ*(*q*)(1 + *F*).

To find the perturbation energy, we use Eq. (15) to write Hamilton’s equation 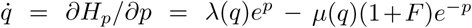, which allows us to find *E*_*p*_ using Eq. (12) upon replacing the lower integration limit by 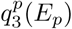. By doing so, we find

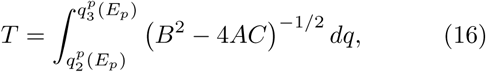

where *A, B* and *C* are given below Eq. (15), and we have swapped the integration limits such that the integrand is positive. Finally, the switching barrier from the *high* to *low* states is given by Eq. (14) upon replacing the lower integration limit by 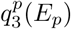, and 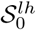 by 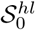.

Fig. 2b shows a comparison of the perturbed *high* to *low* switching probabilities from Monte Carlo simulations with *𝒫*^*hl*^ calculated using the above action along with Eq. (8). They are again in excellent agreement.

### Inference of the epigenetic landscape

We now proceed to our main idea which is to use our theoretical formalism to infer the epigenetic landscape of a regulatory network, given experimental data of the network’s response to external perturbations. The goal is to find the set of parameters for the regulatory network that best recapitulate the observed responses. We first set out to determine the feasibility of inferring the parameters from the perturbation data.

For the SRG, two key parameters that control the shape of the landscape are *β*, which influences the barrier position, and *h*, which influences the landscape steepness. We desired to know to what extent these two parameters could be independently distinguished using only the switching probability. To this end, we used our theory to calculate the dependence of *𝒫*^*lh*^ and *𝒫*^*hl*^ on *β* and *h*. As can be seen in Fig. 2c+d, when switching either from *low* to *high* or from *high* to *low, β* and *h* can be changed simultaneously to maintain the same switching probability. This corresponds, *e.g.*, to moving the barrier position closer to the starting state while increasing the height of the barrier. However, by considering switching in *both* directions simultaneously, both *β* and *h* are uniquely constrained. There is only one pair of values for *β* and *h* that is consistent with both the *low* to *high* and *high* to *low* pulling.

To perform parameter inference we adopted a *maximum likelihood* approach. To generate synthetic experimental data we performed Monte Carlo simulations of the stochastic process (1), and measured for various values of *F* and *T* the number of realizations *k*, out of *m* total realizations, that switched phenotypes after some designated time. For all of our simulations we used *m* = 1×10^6^. Given the switching probability *𝒫* for each realization, and assuming that the sequence of “experiments” or numerical realizations is independent and identically distributed, the probability *P* (*k*) that exactly *k* realizations out of *m* switch is given by a binomial distribution

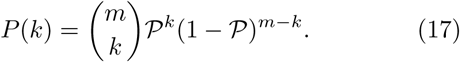

The likelihood of parameters *θ* producing the observed data *k* given all of the various experimental *F* and *T* conditions is then given by the product of all *P* (*k*) values

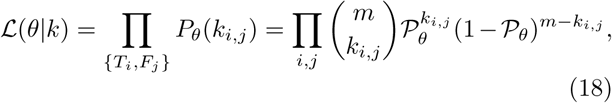

where *i* and *j* denote the indices of the current values of *T* and *F*, and *k*_*i,j*_ denotes the number of realizations that switched given that *T* = *T*_*i*_ and *F* = *F*_*j*_.

Importantly, the probability of success *𝒫*, is given by Eq. (8); as we have shown, it depends, in addition to *T* and *F*, on the parameters *θ* defining the birth and death rates. By maximizing the likelihood function ℒ, we find the most probable parameter set for the birth and death rates Λ_*n*_ and *ℳ* _*n*_, given the perturbation data.

We used the synthetic data set shown in Fig. 2a+b along with Eq.(18) to infer the maximum likelihood estimate (MLE) for the three model parameters *N*, *β*, and *h*. We assume that *α* can be obtained directly from experimental measurement of the ratio of the stable fixed points. Again, we used only *F* and *T* values with 1 ×10^−6^ *< 𝒫 <* 1 ×10^−2^, which amounted to ∼35 experimental conditions combined from both switching directions. Fig. 3 and Fig. S8 show the likelihood distribution resulting from the inference. The MLE was *N* = 1554, *β* = 0.5718, and *h* = 3.492, which was in excellent agreement with the true parameter values of *N* = 1500, *β* = 0.5715, and *h* = 3.5.

**Figure 3.**
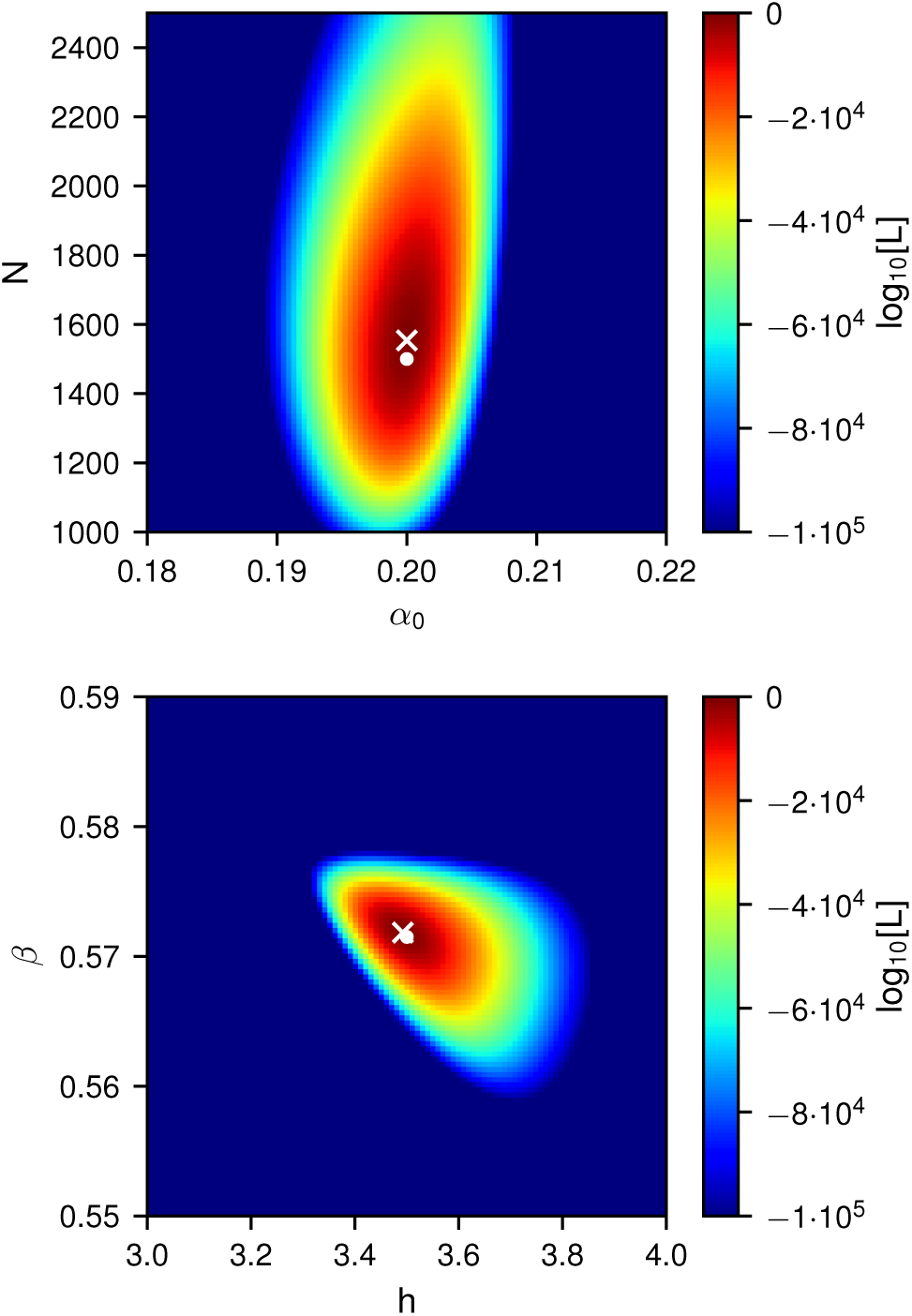
Log-likelihood function for inference of model parameters. (top) The likelihood in the *N* vs *α*0 plane where other parameters are fixed to their MLE. (bottom) The same for the *β* vs *h* plane. The white ×symbols show the MLE and the white ◦ symbols show the true parameter values.

Because the WKB is a logarithmic theory, there is a preexponent in Eq. (8) that must be estimated in order to compute the absolute value of *𝒫*. Typically, in a WKB theory this prefactor is obtained from fitting the functional dependence of the theory to the data. Here, we took the approach of obtaining the prefactor for both *𝒫* ^*lh*^ and *𝒫* ^*hl*^ directly from the likelihood estimation. We maximized likelihood over a range of prefactors and then used the set with the highest likelihood for all remain calculations (see Fig. S4). Comparison of the theory with optimized prefactors and parameters to the true parameters shows that the optimization leads to a moderate increase in likelihood while maintaining the excellent fits to *T* and *F* (see Fig. S12a+b).

Finally, we used the MLE parameters to reconstruct the stationary probability density function (PDF) that corresponds to the epigenetic landscape. Fig. 4a+b shows a comparison of the inferred and true PDFs, which we calculated for a given set of parameters using an enhanced sampling technique [78]. The agreement is again excellent and shows that by using only ∼35 perturbation data points we are able to successfully reconstruct the epigenetic landscape of the model.

**Figure 4.**
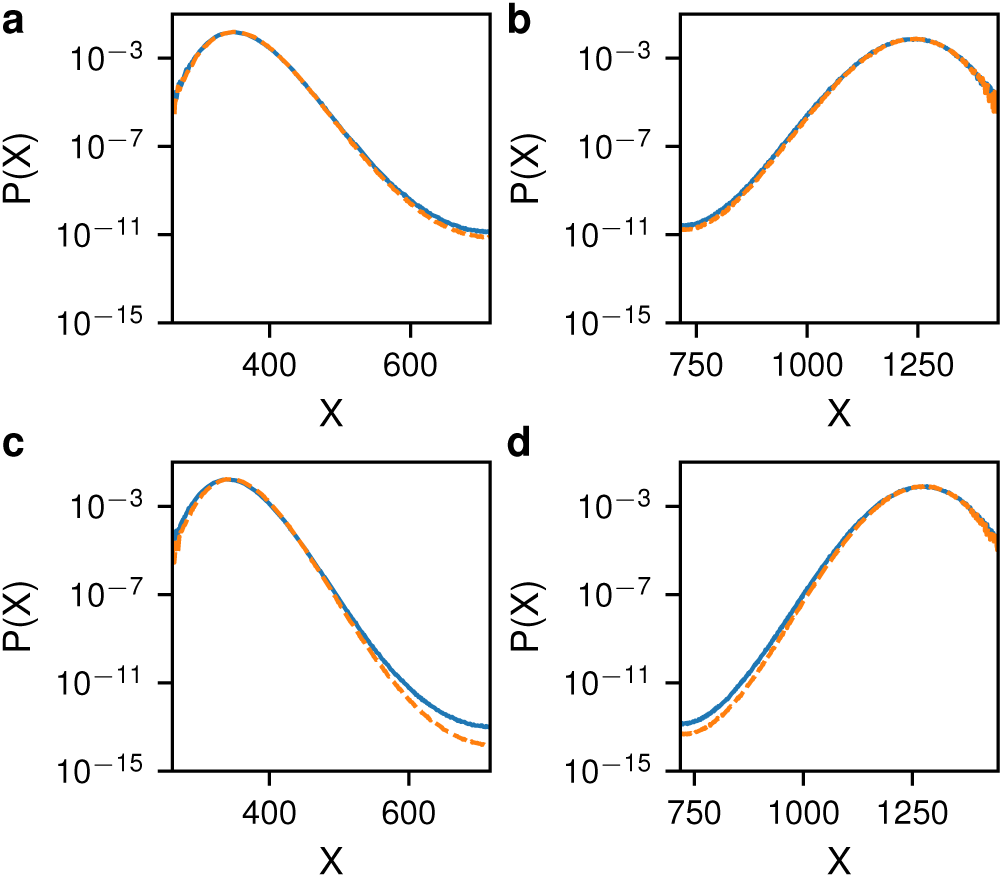
Comparison of actual and inferred probability distributions for the self-regulating gene. (a) The actual (solid blue) and inferred (dashed orange) PDFs for the *low* state. (b) The same for the *hi* state. The model parameters were *N* = 1500, *α*0 = 0.2, *β* = 0.5715 and *h* = 3.5. (c+d) The same as (a+b) except for *h* = 3.65.

To further test our ability to use the theory to infer the network’s PDF, we tested several other SRG parameters sets with increasing switching barrier heights (see Fig. S4–S12). Fig. 4c+d shows that for *h* = 3.65 a greater discrepancy appears between the inferred and actual landscapes, with ∼5% error in the height of the switching barrier. As the switching barrier continues to increase so does the estimated error (Fig. S13). For *h* = 4.0 the error is ∼10% of the barrier height. However, at this value of *h* the MST is ∼ 10^17^. At these very long switching times, emanating from the large landscape steepness, additional perturbation points with *<* 1 ×10^−6^ may be necessary to accurately infer landscapes to within a smaller margin of error.

### One-state mRNA-protein model

#### The unperturbed case

Above, we used the SRG as a basis to infer the landscape of a 1D switch. To see how our method can be generalized to higher-dimensional systems, we now repeat the calculations done above for a 2D system: the one-state mRNA-protein model with positive feedback that displays bistability. We explicitly account for mRNA noise which has been shown to greatly affect the switching properties in genetic circuits [29].

We consider a one-state gene-expression model where transcription depends on the protein copy number via positive feedback. The deterministic rate equation describing the dynamics of the average numbers of mRNA and proteins, respectively denoted by 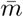 and 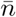, satisfies:

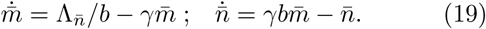

Here, *γ* ≫ 1 is the mRNA degradation rate (relevant *e.g.* for bacterial systems [15]), *γb* is the protein translation rate, such that *b* is the burst size (the number of proteins created from a single instance of mRNA) and all rates are rescaled by the protein’s degradation rate or cell division rate. Furthermore, 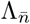 is a sigmoid-like function that ensures bistability (see Fig. S14). By choosing the mRNA transcription rate to be 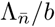, we made sure that the fixed points of the proteins satisfy 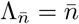, which coincide with those of Eq. (2) for the SRG, upon choosing 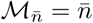.

To find the switching probability we write down the master equation describing the dynamics of *P*_*m,n*_ – the probability to find *m* mRNA molecules and *n* proteins:

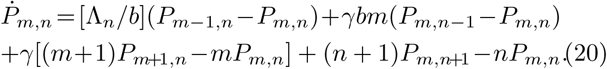

We now go along the same lines as in the case of the SRG. That is, we use the metastable ansatz *P*_*m,n*_ = *π*_*m,n*_*e*^−*t/τ*^ in Eq. (20), and employ the WKB approximation, *π*_*m,n*_ = *π*(*x, y*) = *e*^−*NS*(*x,y*)^, where *S*(*x, y*) is the action, *N* ≫ 1 is the typical protein population size at the *high* state, and *x* = *m/N* and *y* = *n/N* are the mRNA and protein concentrations, respectively. This yields a stationary Hamilton-Jacobi equation *H*(*x, y, ∂*_*x*_*S, ∂*_*y*_*S*) = 0 with Hamiltonian [69, 79]

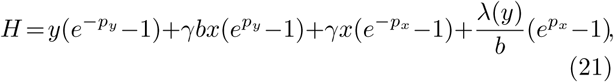

where *λ*(*y*) = Λ(*y*)*/N*, and *p*_*x*_ = *∂S/∂x* and *p*_*y*_ = *∂S/∂y* are the associated momenta of the mRNA and proteins, respectively.

The switching path from the *low* to *high* states (or vice versa) corresponds a heteroclinic trajectory of Hamiltonian (21) connecting the saddle points (*x, y, p*_*x*_, *p*_*y*_) = (*y*_*low*_*/*(*γb*), *y*_*low*_, 0, 0) and (*y*_*high*_*/*(*γb*), *y*_*high*_, 0, 0) in the 4D phase space; it can be found by solving the Hamilton equations 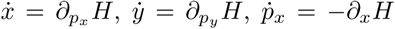 and 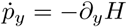, which read

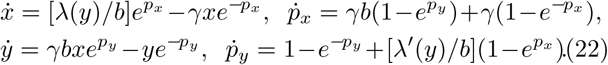

While a numerical solution can be found for any set of parameters, in order to make analytical progress we consider the limit where the mRNA lifetime is short compared to that of the protein, *γ* ≫ 1, which holds in bacteria. In this limit, the mRNA concentration and momentum, *x*(*t*) and *p*_*x*_(*t*), instantaneously equilibrate to some (slowly varying) functions of *y* and *p*_*y*_ [80]. Putting 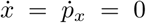 in the first two of Eqs. (22), we obtain 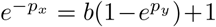 and 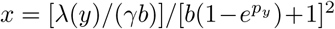. Using these relations in Hamiltonian (21) we arrive at *reduced* Hamiltonian for *y* and *p*_*y*_ only. Denoting *q* ≡ *y* and *p* ≡ *p*_*y*_, the effective 1D Hamiltonian reads [69, 79]

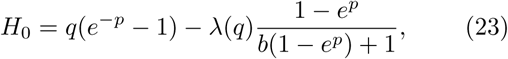

where the subscript 0 denotes the unperturbed case. This Hamiltonian effectively accounts for the fact that the proteins are produced in geometrically distributed bursts with mean *b*, which in turn asymptotically accounts for the mRNA noise when *γ* ≫ 1. This Hamiltonian is our starting point for treating this system under external perturbation, and serves as the unperturbed Hamiltonian, similarly as Hamiltonian (5). Note that, as done for the SRG model above, using this unperturbed Hamiltonian [Eq. (23)], one can find the unperturbed action which yields the PDF *P*_*m,n*_ and MST, in the absence of external perturbation, see Fig. S15+S16,.

#### The perturbed case

Now we repeat the calculations done for the SRG in the perturbed case. Note, that here, instead of perturbing the protein’s expression and degradation rates, we perturb those of the mRNA. While both cases can be studied theoretically, we desired to study the impact of transcriptional perturbations as being more closely aligned with existing experimental techniques.

We start by perturbing the mRNA’s transcription rate Λ_*n*_ →Λ_*n*_ + *ϕ*(*t*), where *ϕ*(*t*) is given by Eq. (9). As before, the optimal path is made of three segments: an unperturbed segment before the onset of perturbation, a perturbed segment while the perturbation is applied, and an unperturbed segment after the perturbation has terminated. The perturbed segment is found by equating Hamiltonian (23) to zero

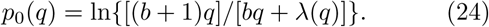

The perturbed segment can be found by using Hamiltonian (23) with the perturbed transcription rate

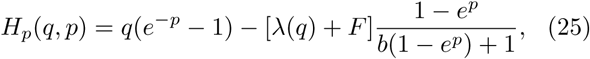

and equating it to *E*_*p*_; here the subscript *p* stands for perturbation. The resulting perturbed segment reads

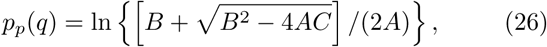

where *A* = *λ*(*q*) + *F* + *b*(*E*_*p*_ + *q*), *B* = *E*_*p*_(*b* + 1) + *λ*(*q*) + *F* + *q*(1 + 2*b*), and *C* = (1 + *b*)*q*. To determine the energy *E*_*p*_, we use Eq. (12) with 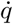 found from Hamiltonian (25). By doing so, condition (12) becomes:

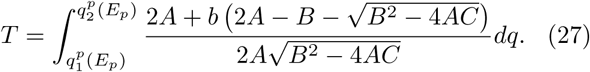

Finally, the action is given by Eq. (14) with *E*_*p*_ from Eq. (27), while 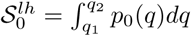 is the unperturbed action from the *low* to *high* states.

Next, we perturb the degradation rate of the mRNA such that *γm* becomes *γ*(1+*F*)*m*. As a result, after some algebra the perturbed Hamiltonian becomes

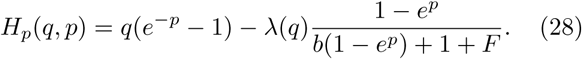

As a result, the perturbed segment satisfying the equation *H*_*p*_(*q, p*) = *E*_*p*_ reads

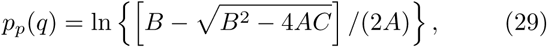

where *A* = *λ*(*q*) + *b*(*E*_*p*_ + *q*), *B* = *E*_*p*_(1 + *b* + *F*) + *λ*(*q*) + *q*(1 + 2*b* + *F*), and *C* = *q*(1 + *b* + *F*). To determine the energy *E*_*p*_, we use Eq. (12) with 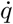 found from Hamilto-nian (28). By doing so, condition (12) becomes:

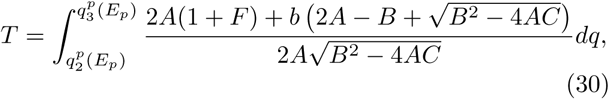

where we have swapped the integration limits such that the integrand is positive. Finally, the action is given by Eq. (14) with E*p* from equation (30), while 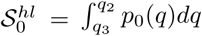 is the unperturbed action from the *high* to *low* states.

To test the mRNA-protein model theory, we again ran sets of Monte Carlo simulations to calculate the dependence of *P*^*lh*^ and *P*^*hl*^ on F for five values of T with b = 4. We then performed MLE estimation from these data, see Fig. S18+S23. Fig. 5a+b shows that there was an excellent agreement between the simulations and the theory with the MLE parameters. Likewise, the reconstructed PDFs shown in Fig. 6a+b are in good agreement with the actual PDFs.

**Figure 5.**
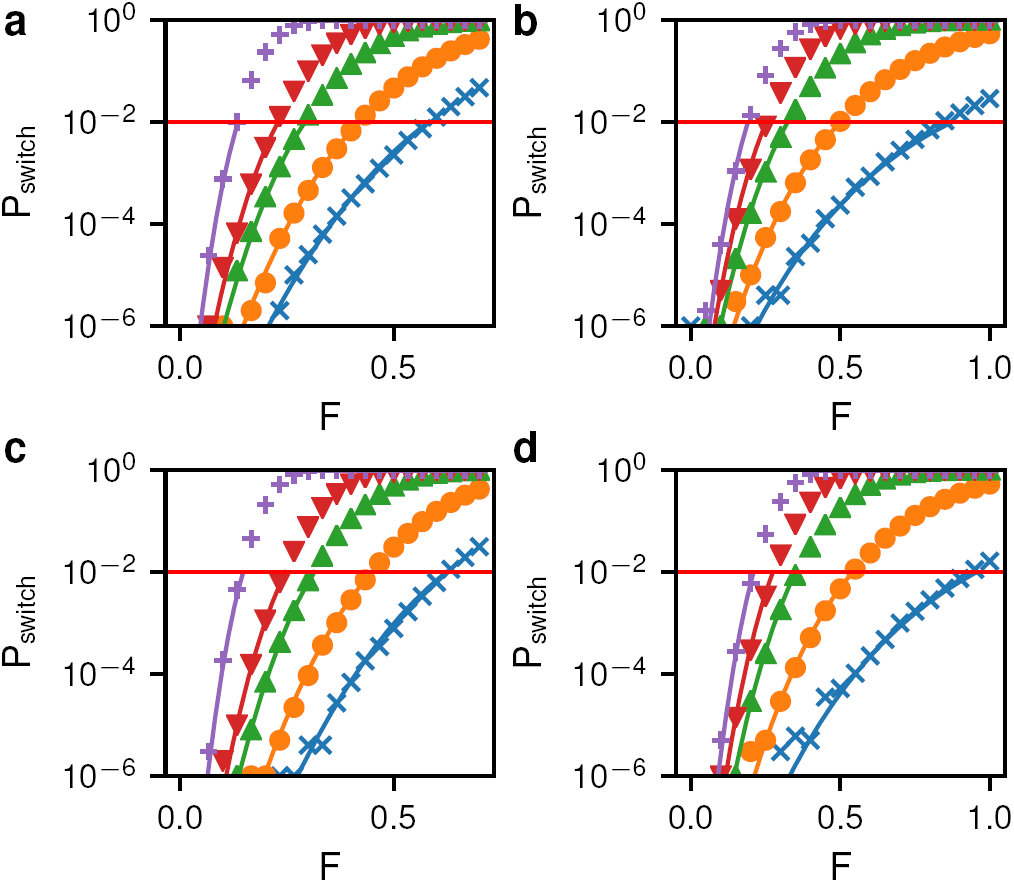
Perturbation effect on the mrna-protein model. (a) Change in switching probability vs perturbation strength *F* for the *low* to *high* switch with *b* = 4. Symbols and lines give numerical and theoretical values, respectively. Theoretical values were calculated using the maximum likelihood parameter estimates. Data are shown for five values of the perturbation time *T* : 0.35 (blue ×), 0.5 (orange ◦), 0.75 (green Δ), 1.0 (red ▽), 2.0 (purple +). (b) Change in switching probability vs perturbation strength for the *high* to *low* switch with perturbation times 1.0 (blue ×), 1.5 (orange ◦), 2.25 (green Δ), 3.0 (red ▽), 4.5 (purple +). (c+d) As above except for *b* = 3.

**Figure 6.**
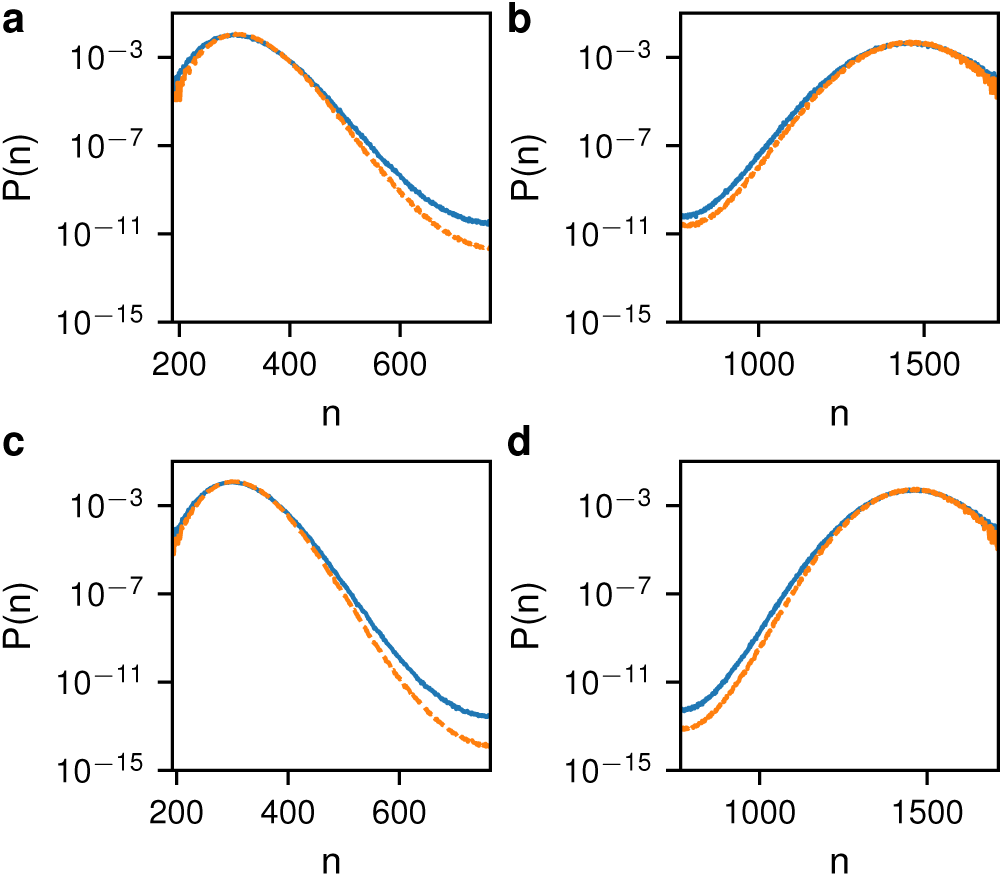
Comparison of actual and inferred probability distributions for the mrna-protein model. (a) The actual (solid blue) and inferred (dashed orange) PDFs for the *low* state with *b* = 4. (b) The same for the *hi* state. (c+d) The same as (a+b) except for *b* = 3.

Finally, we wanted to study the impact of increasing the barrier height while maintaining the position of the fixed points. To this end, we varied *b* in a range of 1–5 (Fig. S17-S28). For parameter sets with longer switching times the low *F* region is not well sampled (Fig. 5c+d), which leads to an increased error in the predicted switching barrier (Fig. 6c+d). With *b* = 1 and a switching time of 1 10^20^ the relative error in the barrier height is ~20%. As with the SRG, these errors could be reduced by including lower probability events in the maximum likelihood estimation.

## Discussion

### Generic models

In general, we would like to apply our methodology to arbitrarily complex networks, not only the simple models discussed above. Let us consider a generic gene-regulatory network with *M* species, **n** = *n*_1_, *n*_2_, …, *n*_*M*_, describing *e.g., M* different proteins that affect each other. Currently, inferring the *M* -dimensional epigenetic landscape in this case could only be done by simulating the dynamics with given reactions. However, our method allows one to find an effective epigenetic landscape, as we did for the mRNA-protein one-state system.

To construct an effective landscape, one needs to find an *order parameter ω*, which is a function of **n** that qualitatively captures the switching dynamics of the *M* species; that is, this order parameter must monotonically change along the system’s switching path. For example, in a genetic toggle switch with protein *A* inhibiting protein *B* and *vice versa*, an adequate choice for an order parameter would be *ω* = (*A − B*)*/*(*A* + *B*). This variable, which fluctuates between 1 and 1 and vice versa, monotonically changes along both switching paths of the system, and thereby effectively captures the switching dynamics of the 2D system [81–83].

Let us assume that in a *M* -dimensional switch, we have found an order parameter *ω* such that its deterministic bistable dynamics are given by

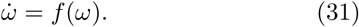

Here *f* (*ω*) = −(*ω* − *ω*_1_)(*ω* − *ω*_2_)(*ω* − *ω*_3_), such that *ω*_*1*_*< ω*_*2*_*< ω*_*3*_ are the fixed points of this equation, Where *ω*_1_ and *ω*_3_ are attracting, while *ω*_2_ is repelling. Note, that time can always be rescaled to have such a bistable dynamics.

To account for noise, we can write a Langevin equation for *ω* satisfying:

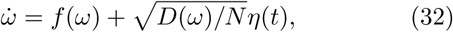

where *D*(*ω*) is an *ω*-dependent diffusion coefficient, *N* ≫ 1 is the typical system size, while *η*(*t*) is a delta-correlated normal random variable with mean 0 and variance 1*/dt*. For concreteness one can choose a cubic function for *D*(*ω*) such that *D*(*ω*) = *D*_0_ + *D*_1_*ω* + *D*_2_*ω*^2^ + *D*_3_*ω*^3^. Under the Itô calculus, this Langevin equation corresponds to the following Fokker-Planck equation for *P* (*ω, t*) – the probability to find order parameter *ω* at time *t*:

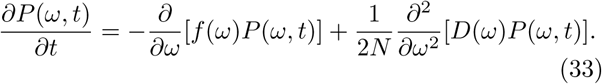

The zero-current steady-state solution (assuming an exponentially small current through the unstable fixed point) of this equation satisfies

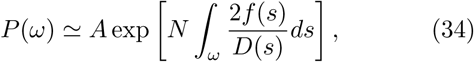

where *A* is a normalization constant, such that 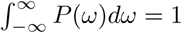.

In analogy to the 1D treatment that we have presented in this paper, we can use the WKB method to infer the epigenetic landscape by perturbing the system with some temporary perturbation of magnitude *F* and duration *T*. This perturbation can be done in any of the proteins that comprise the system. Indeed, plugging the WKB ansatz 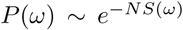 into Eq. (33), in the leading order in *N* ≫ 1 one arrives at a Hamilton-Jacobi equation with a Hamiltonian satisfying:

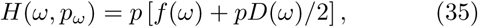

where *p* = *dS/dω* is the conjugate momentum. As a result, the unperturbed optimal path, *p*_0_(*ω*), satisfies

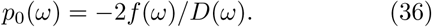

At this point we can repeat the analysis done for the models above. By writing down the perturbed Hamiltonian, *H*_*p*_, and equating it to *E*_*p*_, one can find the perturbed trajectory segment, *p*_*p*_(*q*), whose intersection points with *p*_0_(*q*) determine *E*_*p*_, by demanding that the perturbation duration be *T*, see Eq. (12). Once these have been computed, we can find the correction to the switching probability as a function of *F* and *T*, according to Eq. (14), both when starting from the vicinity of *ω* = *ω*_1_ or *ω* = *ω*_3_.

Once the theoretical results are available, one can employ the maximum likelihood method to infer the parameters in this model that determine the deterministic drift function *f* (*ω*) and the *ω*-dependent diffusion *D*(*ω*). We envision that this may lead to a general method for studying epigenetic landscapes using perturbations.

#### Dependence of switching probability on impulse

In physical terms *FT* represents the total *impulse* we apply to the system, which is equal to the force exerted on a particle multiplied by the duration of the force. In a mechanical system, when a constant force *F* is applied on a particle for a duration *T* in the direction of the particle’s momentum, in the absence of dissipation or heat production, the particle’s momentum is increased by *FT*. This increase is independent on *F* or *T* separately; that is, applying a small force for a long duration is equivalent to applying a large force for a short duration.

This relationship is exactly what we observe for small impulses in our system. Fig. 7 shows the change in the switching barrier *e.g.* between the *low* and *high* states, 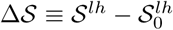 as a function of total impulse *FT*. One can see that for low *FT* the change in the switching barrier depends linearly on the product *FT* and not on *F* or *T* separately. However, as the impulse increases the change in switching barrier is no longer a unique function of *FT*. As the impulse duration *T* is increased (high *FT* with low *F*) not all of the impulse results in a reduction in the switching barrier. This discrepancy indicates that there is some sort of dissipation or uncontrolled heat/entropy production in the system.

**Figure 7.**
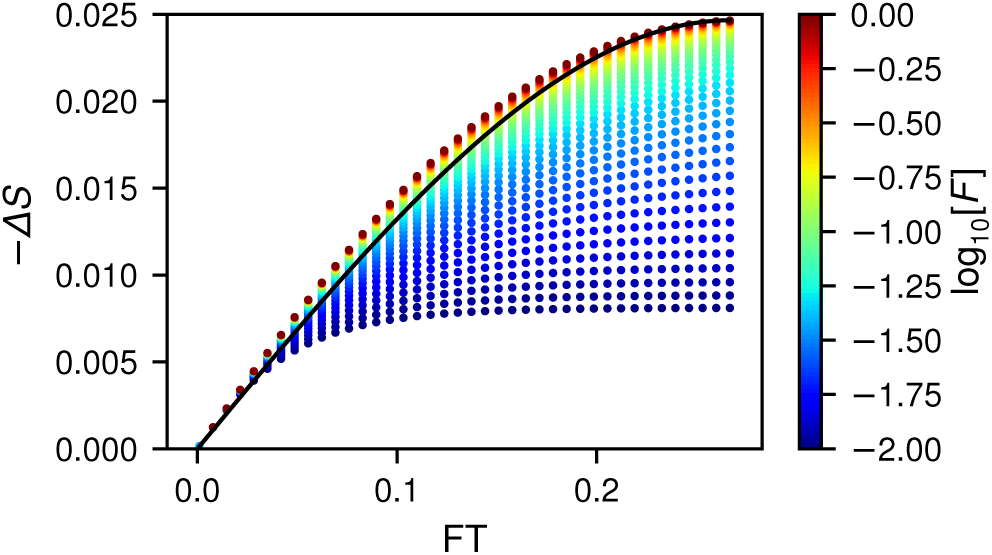
Dependence of change in the switching barrier on impulse. Distribution of the perturbed switching barrier 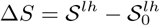 versus the total applied impulse *F T* for the SRG model. Each symbol represents a perturbation with a different *F*, which is given by the color. The solid line shows the bifurcation theory given in Eq. (44).

In contrast, when the system is near bifurcation we do expect the change in action to be a unique function of *FT*. In Appendix A we derive a simple analytical expression for the dependence of the switching barrier on *FT* close to the bifurcation limit. As can be seen in Fig. S29, the effect of the perturbation depends only on the product *FT* in this case.

Even though the switching barrier is a unique function of *FT* close to bifurcation, see Eq. (44), we can see that the change in switching barrier is linear with *FT* only at low impulse. Defining the *efficiency* of inducing a switch by the change in the switching barrier divided by the impulse, Δ*S/*(*FT*), from Eq. (44) we see that the efficiency decreases with *FT*. That is, the process of inducing a switch becomes less efficient as *FT* is increased, while efficiency is maximized for weak impulses with vanishingly-small dissipation.

#### Role of perturbation energy E_p_

What is the physical meaning of the perturbation energy *E*_*p*_ which appears throughout our derivation? Mathematically, it is determined by a complicated function of the force *F* and its duration *T*. However, looking at the result close to bifurcation (see Appendix A), the energy *E*_*p*_ can be written as *E*_*p*_ = *E*_0_[1 − (*FT*)^2^*/*4], where *E*_0_ is the maximal value of *E*_*p*_ which is obtained as *F* and/or T vanish impulses 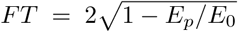. This indicates that, in analogy to quantum mechanics, given a quasi-potential *T* vanish. Plugging this result into Eq. (44), and sub-stituting 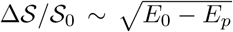, we find that for small *S* ≡ *S – S* landscape *S*(*q*), *E*_0_ − *E*_*p*_ can be viewed as the “energy excess” the particle receives to cross the switching barrier of height *E*_0_. As a result, for *E*_*p*_ = *E*_0_, the switching barrier remains unchanged (corresponding to *F* = 0 and/or *T* = 0), while for *E*_*p*_ = 0, the energy excess is maximal corresponding to the absence of a switching barrier, leading to instantaneous switching.

## Conclusion

Here, we have introduced a new theory for describing the effect of nonequilibrium perturbations on biological regulatory networks with metastable states, *i.e.*, epigenetic networks. Our theory can be used to infer the epigenetic landscape of a regulatory network by fitting a model using a series of perturbations of varying strength. The shape of the landscape is mapped out and reconstructed from the perturbation responses. The data needed for the fitting are purposely chosen to be reasonable biological observables.

The principle of such an experiment would be to apply a genetic or biochemical perturbation to the network, such as by introducing an inducible gene using transfection and/or silencing expression using siRNA. A response is measured as a function of the strength and duration of the perturbation. Unlike other theories that relate switching dynamics to fluctuations along the epigenetic landscape, our theory does not require detailed time-lapse imaging to collect data. One simply needs to record the fraction of cells that switch phenotypes at some time in the future after the perturbation. Such data can be quickly collected for millions of cells using flow cytometry. Using the response data the parameters of a regulatory model can be inferred and the epigenetic landscape numerically reconstructed. Our method, therefore, has great potential to be used to help decipher complex biological developmental trajectories.

Our theory also provides insights into the fundamental physics of how various signals induce state transitions in cells. As more complex cellular reprogramming is undertaken, it will become increasingly important to model how cells can be induced to make transitions between states. The impulse that we identified in our theory is one way to measure the work required to change phenotypic states. Further theoretical advances will be required to extend our understanding of important developmental techniques such as creation of induced pluripotent stem cells and cell reprogramming and differentiation.

## Acknowledgments

The authors would like to thank Chris Bohrer and Tom Israeli for early discussions on the theoretical framework. ER acknowledges support from the National Science Foundation under grant PHY-1707961. MA and SB acknowledge support from the Israel Science Foundation grant No. 300/14 and the United States-Israel Binational Science Foundation grant No. 2016-655.

## Appendix A: Bifurcation Limit

In this section we show how the results derived for the SRG can be drastically simplified close to the bifurcation limit, where the stable and unstable fixed points merge. Without loss of generality, our analysis below will focus on the case of switching from the *low* to *high* states, namely from *q*_1_ to *q*_3_.

Let us denote by *ϵ* ≡ (*q*_2_−*q*_1_)*/*2 ≪ 1, such that 2*E* is the distance between the two fixed points. Let us also denote by *q*_*m*_ = (*q*_1_ + *q*_2_)*/*2 the mid point between the stable and unstable fixed points. As a result, we can write *q*_1_ = *q*_*m*_ −*ϵ* and *q*_2_ = *q*_*m*_ + *ϵ*. It has been shown in previous works that in problems of switching between metastable states, close to bifurcation the momentum *p* scales as *ϵ*^2^ [65, 66]. As a result, it is convenient to rescale the coordinate and momentum as follows:

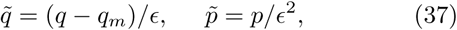

where 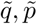 are *𝒪* (1). Note, that the fixed points in the rescaled coordinate become 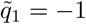 and 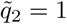.

We further denote *R*(*q*) = *λ*(*q*)−*µ*(*q*), such that in the absence of external forcing the mean-field rate equation becomes 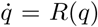. Since *R*(*q*) can be approximated by a parabola in the regime *q*_1_ *< q < q*_2_, we have: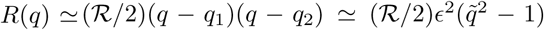, where R≡*R*″(*q*_*m*_) is a positive constant. Therefore, at the midpoint *R*(*q* = *q*_*m*_) = −(*R/*2) *ϵ*^2^ is negative, and *R*′(*q*_*m*_) = 0, since the parabola has a minimum at *q* = *q*_*m*_. Using these results, and denoting by *𝒟* ≡ *λ*(*q*_*m*_) + *µ*(*q*_*m*_), we now expand the time-dependent Hamiltonian up to *𝒪* (*ϵ*^4^) in the vicinity of *q* = *q*_*m*_ and *p* = 0. This results in

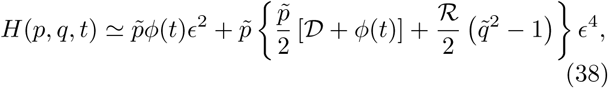

where we have expanded 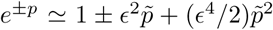, and have also expanded *λ*(*q*) and *µ*(*q*) around *q* = *q*_*m*_, up to second order in *ε*.

In the absence of external force, *ϕ*(*t*) = 0, the unperturbed optimal path satisfies *H* = 0, which yields the unperturbed trajectory

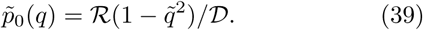

In the presence of an external force, *ϕ*(*t*) = *F*, the perturbed Hamiltonian becomes in the leading order

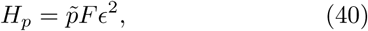

independent of 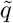. Thus, equating *H*_*p*_ = *E*_*p*_ yields the perturbed optimal path, which becomes constant here

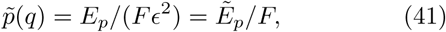

where we have defined the rescaled energy 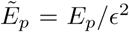. Equating the non-perturbed and the perturbed optimal paths, Eqs. (39) and (41), we find the intersection points 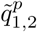 to be 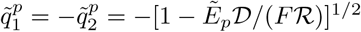.

Let us now find the rescaled energy given the duration of the external perturbation *T*. Using Eqs. (12) and (40), and the fact that 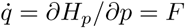, we have

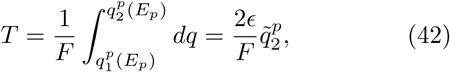

from which we can extract *E*_*p*_ as a function of *T*:

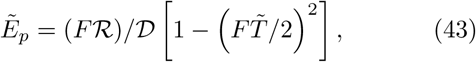

where 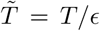. Note, that the result is valid as long as 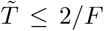, which means that *T≤*2 *∊/F*. The is because when the system is close to bifurcation, a very small force, *F ∽ ∊* is sufficient to cause a deterministic switch. Therefore, if *F ∽ ∊*, we have *T* = 𝒪 (1). Also note that, by using Eq. (43), the intersection points become 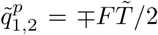; here, at the maximal value of 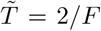. we obtain *E*_*p*_ = 0, since 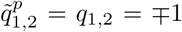. coincide with the unperturbed fixed points *q*_1,2_.

Having found the perturbation energy, the correction to the switching barrier is given by Eq. (14). Transforming to the rescaled coordinate and momentum, using Eqs. (39), (41) and (43), and using the definition of 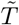, we finally have

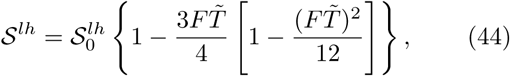

where 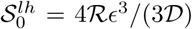, and the result is valid as long as 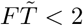. Fig. S29 shows a comparison of Eq. (44) and the full theory when the system is near bifurcation. Note, that as *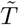* approaches 2*/F*, action (44) approaches zero, which invalidates the WKB theory. The latter is valid as long as *Ns*≫ 1, which limits the duration and/or magnitude of the external force.

## Appendix B: Weak noise limit

In this section we derive the switching barrier under a weak external perturbation. Here, one must have a long perturbation duration; otherwise the effect is negligible. In the following we will assume for simplicity that *E*_*p*_ = 0 which corresponds to a long perturbation duration, see below.

When *E*_*p*_ = 0, the perturbed and unperturbed optimal paths for switching intersect at *q*_1_ and *q*_2_, such that 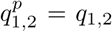. Therefore, taking Eq. (11), putting *E*_*p*_ = 0 and expanding in *F* ≪ 1, the perturbed optimal path becomes

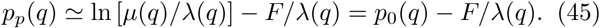

As a result, using Eq. (14), and the fact that *E*_*p*_ ≃ 0, the correction to the switching barrier in the case of weak force, drastically simplifies and becomes

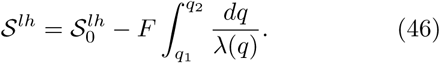

Note, that in this case, the duration of the external perturbation is simply given by 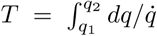, where 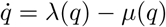 is the unperturbed rate equation.

## Appendix C: Heaviside Step Function Translation Rate

In this section we consider the case of very large Hill exponent in Eq. (3). In the limit *h → ∞* the translation rate becomes a step function

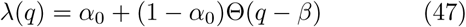

where *α*_0_ = *a*_0_*/N*, *β* = *n*_0_*/N* and Θ(*z*) is a heaviside step function. In this case, the mean-field rate equation has three fixed points: *q*_1_ = *α*_0_, *q*_2_ = *β* and *q*_3_ = 1, where *q*_1_ and *q*_3_ are stable, while *q*_2_ is unstable.

Let us begin by computing the correction to the switching barrier from the *low* to *high* states. Here the un-perturbed switching barrier satisfies 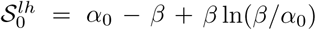 [33]. Going along the same lines as above, we can compute the unperturbed and perturbed optimal paths for switching using Eqs. (6) and (11), as well as the intersection points between these paths, which satisfy 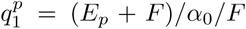 and 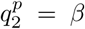. In this case, one can explicitly find *E*_*p*_ using the expression for 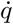 and Eq. (13). The result is

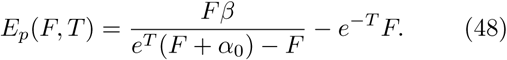

Using this result and Eq. (14), the correction to the switching barrier from the *low* to *high* states is:

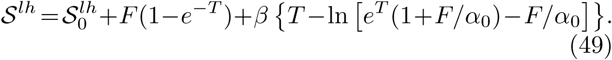

We now move to compute the correction to the switching barrier from the *high* to *low* states. Here the un-perturbed switching barrier satisfies 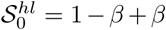 In *β*. Going along the same lines as above, we can compute the unperturbed and perturbed optimal paths for switching using Eqs. (6) and (11) as well as the intersection points between these paths, which satisfy 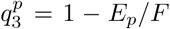 and 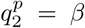. In this case, one can also explicitly find *E*_*p*_ using the expression for 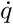 and Eq. (12) with the lower integration limit replaced by 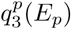. The result is

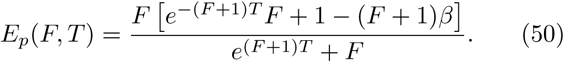

Using this result and Eq. (14), the correction to the switching barrier from the *high* to *low* states is:

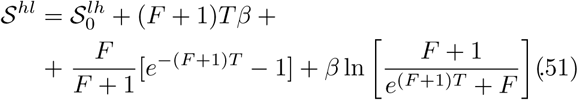

