## Supplementary Material for "Reconstructing an epigenetic landscape using a genetic ‘pulling’ approach"

### Supplementary Material for
### "Inferring the epigenetic landscape using a genetic 'pulling' approach"

Michael Assaf[1⋆], Shay Be'er[1] and Elijah Roberts[2†]

September 20, 2019

[1] Racah Institute of Physics, Hebrew University of Jerusalem, Jerusalem 91904, Israel

[2] Department of Biophysics, Johns Hopkins University, Baltimore, MD 21218, USA

⋆ Correspondence to:

Michael Assaf

Racah Institute of Physics, Hebrew University of Jerusalem

Popick 116

Givat Ram

Jerusalem, Israel 91904

Ph: +972-2-6584593

† Correspondence to:

Elijah Roberts

Department of Biophysics, Johns Hopkins University

Jenkins Hall 110

3400 N Charles St

Baltimore, MD 21218

Ph: +1-410-516-2384

#### Supplementary Figures

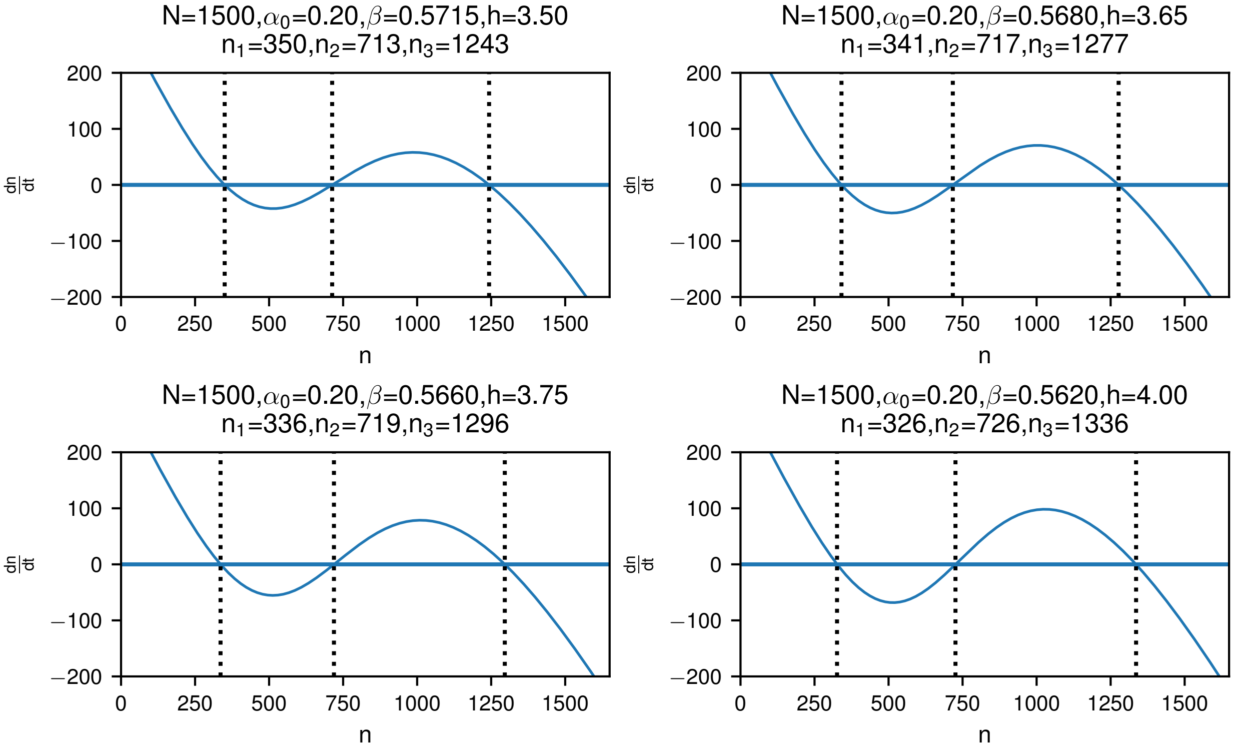


**Figure S1**: **Deterministic rate equations for the self-regulated gene model.** Value of $\frac{dn}{dt}$ as a function of $n$ for the deterministic model of the SRG. The positions of the three fixed points are given by the dotted lines. Parameters for each panel are as indicated.

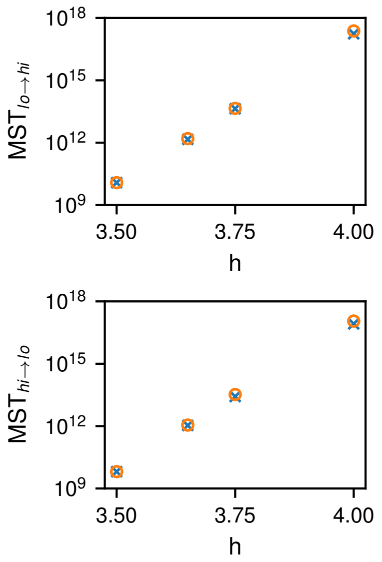


**Figure S2**: **Mean switching times for the self-regulated gene model.** (top) The MST to go from the $low$ state to the $high$ state vs $h$ calculated from numerical simulations (blue $\times$) and WKB theory (orange $\circ$) as given by Eq. (7) in the main text. The WKB points are multiplied by a constant preexponential factor of 20.4. (bottom) The same for the $high$ to $low$ state with a preexponent of 11.94. All other parameters are as in Figure S1.

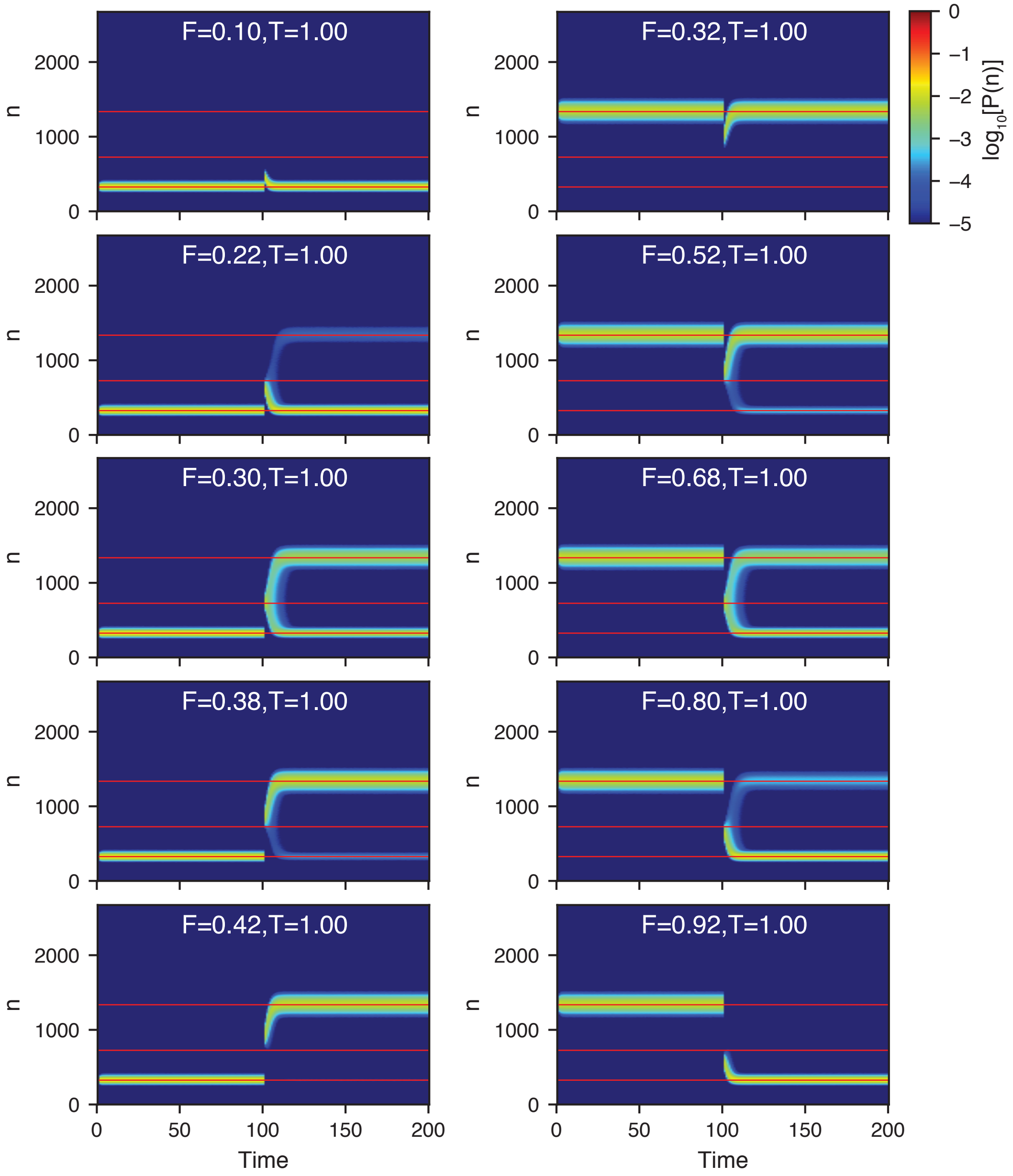


**Figure S3**: **Time dependent probability distribution for pulling on the self-regulated gene model.** The plots show the probability distribution versus time for pulling from $low \to high$ (left column) and $high \to low$ (right column). The simulations were initialized at the appropriate fixed point and allowed to equilibrate until t=100.0 before pulling was initiated. A pulling force of strength F was applied for the specified time T and then removed. The simulations then continued running until t=200.0 to relax and then the switching statistics were calculated. Red lines show the locations of the deterministic fixed points. Parameters for the SRG model were $N = 1500$, $\alpha_0 = 0.2$, $\beta = 0.562$, and $h = 4.0$.

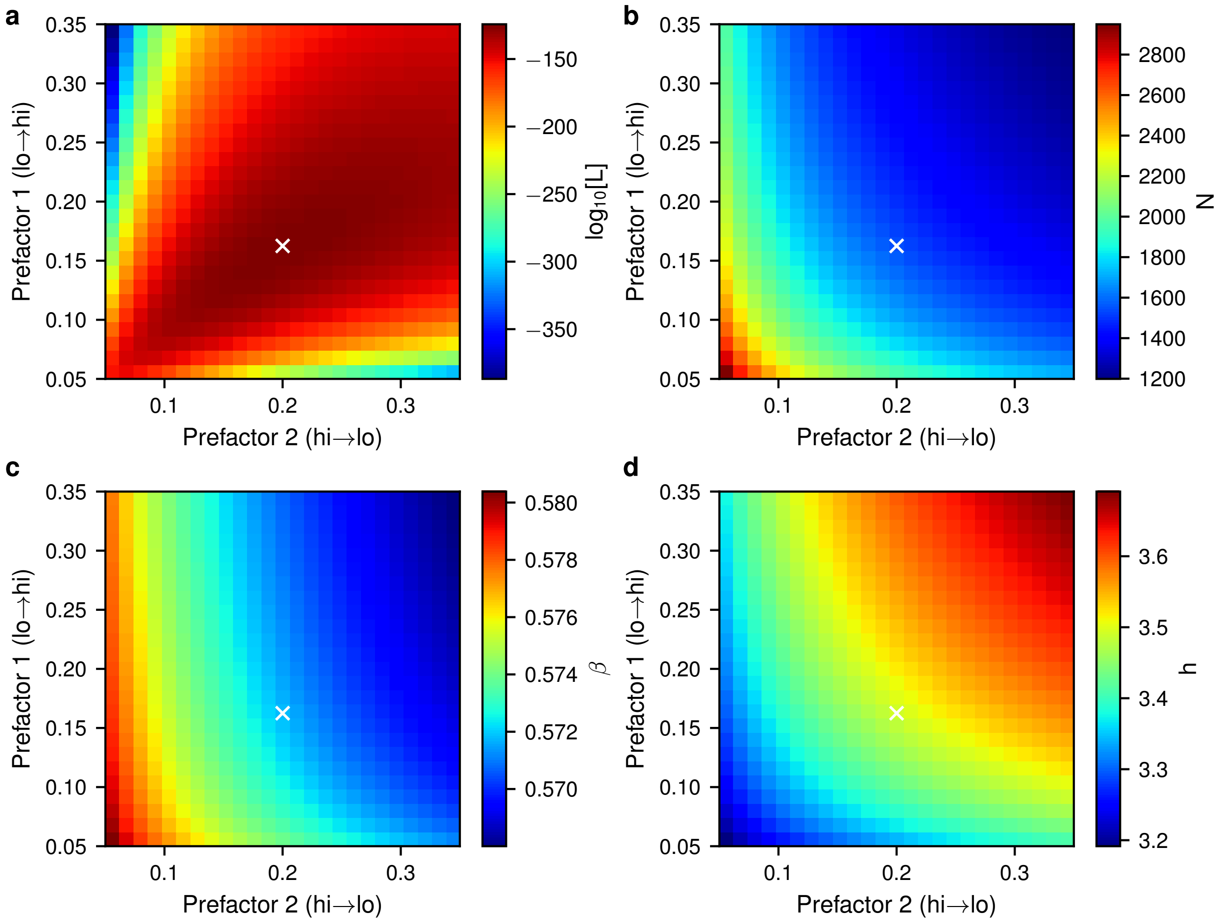


**Figure S4**: **Prefactor dependence during maximum likelihood fitting of the SRG model with $h = 3.5$.** (a) The maximum likelihood score obtained for each prefactor pair during fitting. Fitting was performed with $\alpha$ fixed to its true value, as described in the main text, using simulation data obtained from the SRG model with parameters $N = 1500$, $\alpha_0 = 0.2$, $\beta = 0.5715$, $h = 3.5$. The prefactor pair with the highest likelihood score is marked with a white $\times$ in each panel. (b-d) The dependence of the maximum likelihood estimate for the parameters $N$, $\beta$, and $h$, respectively, on the prefactors.

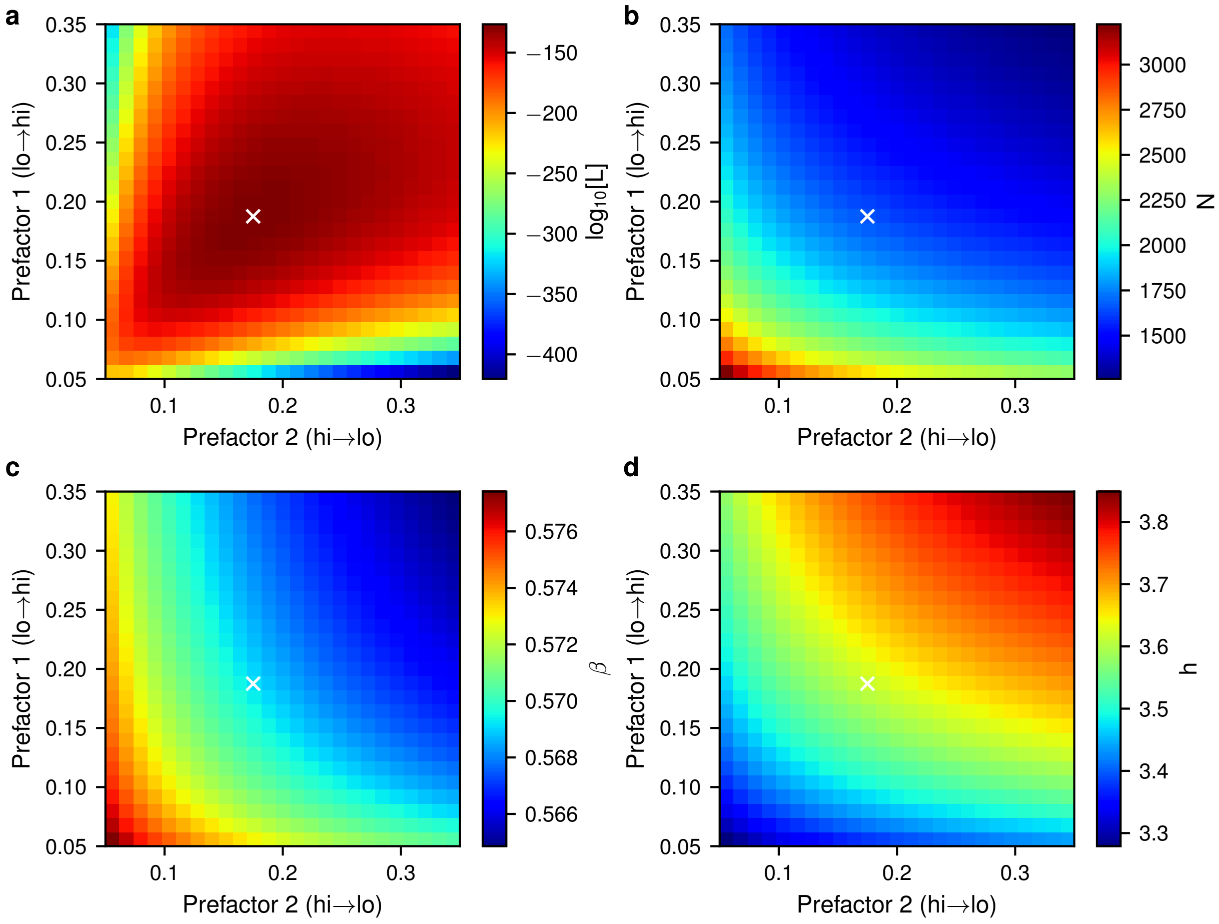


**Figure S5**: **Prefactor dependence during maximum likelihood fitting of the SRG model with $h = 3.65$.** (a) The maximum likelihood score obtained for each prefactor pair during fitting. Fitting was performed with $\alpha$ fixed to its true value, as described in the main text, using simulation data obtained from the SRG model with parameters $N = 1500$, $\alpha_0 = 0.2$, $\beta = 0.568$, $h = 3.65$. The prefactor pair with the highest likelihood score is marked with a white $\times$ in each panel. (b-d) The dependence of the maximum likelihood estimate for the parameters $N$, $\beta$, and $h$, respectively, on the prefactors.

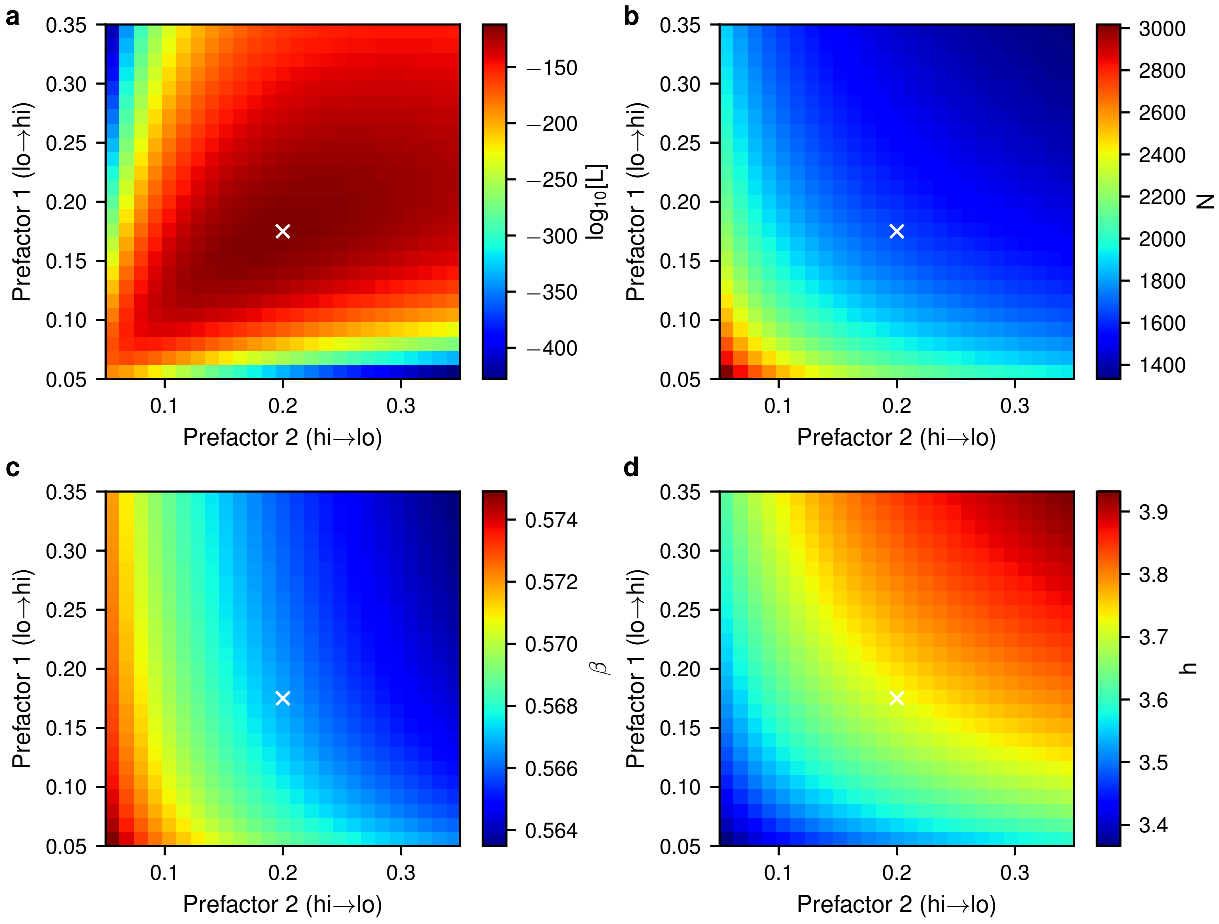


**Figure S6**: **Prefactor dependence during maximum likelihood fitting of the SRG model with $h = 3.75$.** (a) The maximum likelihood score obtained for each prefactor pair during fitting. Fitting was performed with $\alpha$ fixed to its true value, as described in the main text, using simulation data obtained from the SRG model with parameters $N = 1500$, $\alpha_0 = 0.2$, $\beta = 0.566$, $h = 3.75$. The prefactor pair with the highest likelihood score is marked with a white $\times$ in each panel. (b-d) The dependence of the maximum likelihood estimate for the parameters $N$, $\beta$, and $h$, respectively, on the prefactors.

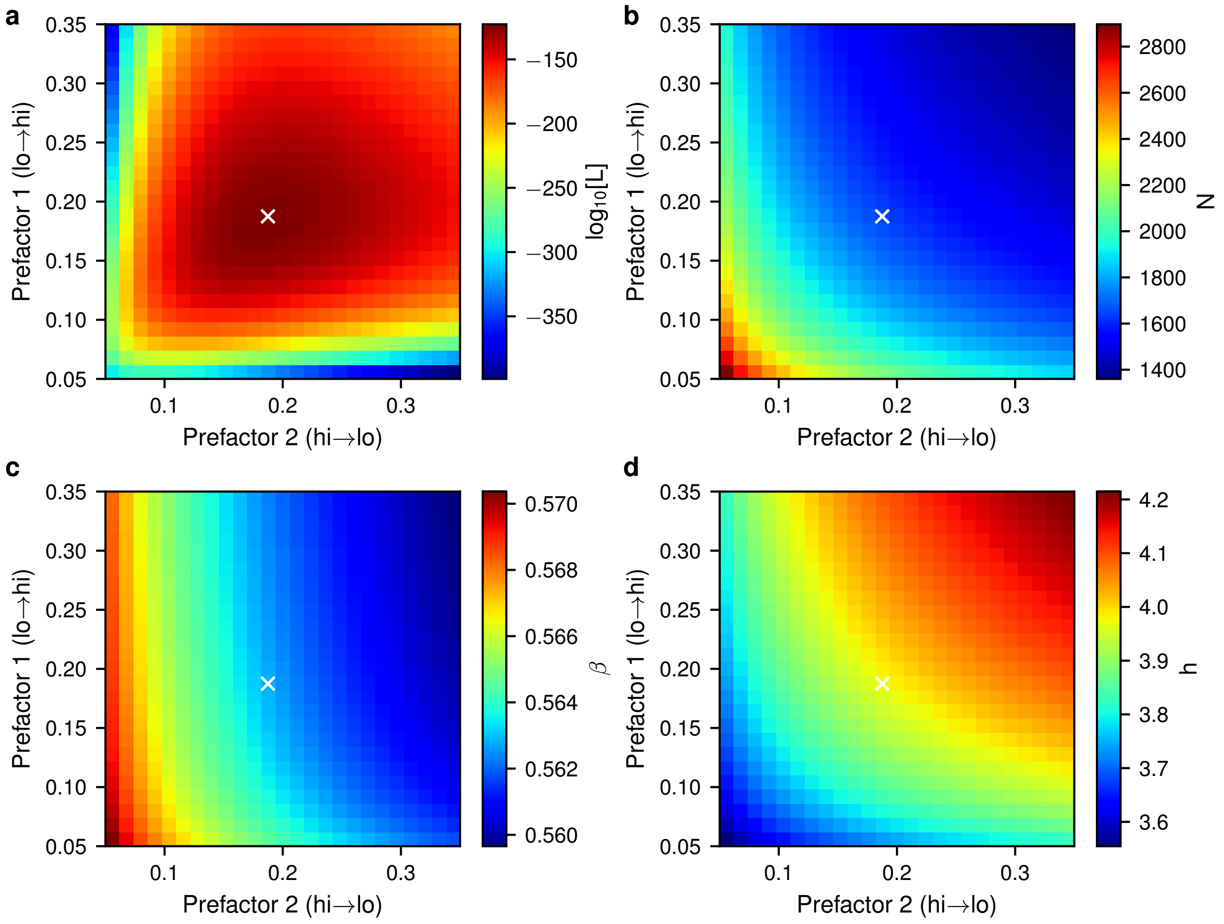


**Figure S7**: **Prefactor dependence during maximum likelihood fitting of the SRG model with $h = 4.0$.** (a) The maximum likelihood score obtained for each prefactor pair during fitting. Fitting was performed with $\alpha$ fixed to its true value, as described in the main text, using simulation data obtained from the SRG model with parameters $N = 1500$, $\alpha_0 = 0.2$, $\beta = 0.562$, $h = 4.0$. The prefactor pair with the highest likelihood score is marked with a white $\times$ in each panel. (b-d) The dependence of the maximum likelihood estimate for the parameters $N$, $\beta$, and $h$, respectively, on the prefactors.

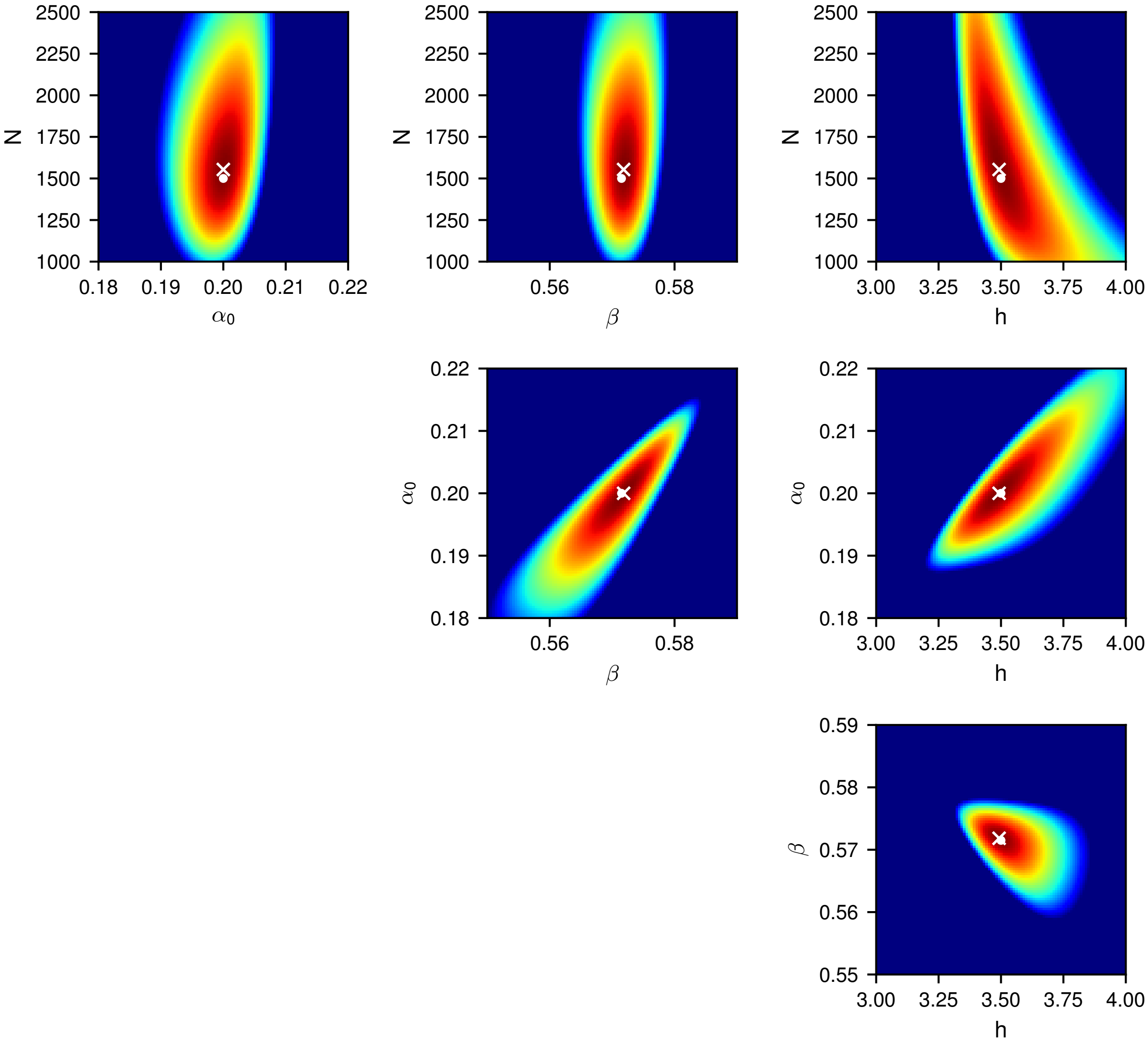


**Figure S8**: **Likelihood distribution for the SRG model with $h = 3.5$.** Likelihood distribution for inference of all pairs of parameters for the SRG model, using the optimized prefactors. For each plot, all other parameters are fixed to their MLE. The MLE is marked with a white $\times$ and the true parameter values are marked with a white •. Colors show $\log_{10}[\mathrm{L}]$ and range from $-1.0{\times}10^5$ (blue) to $0$ (red).

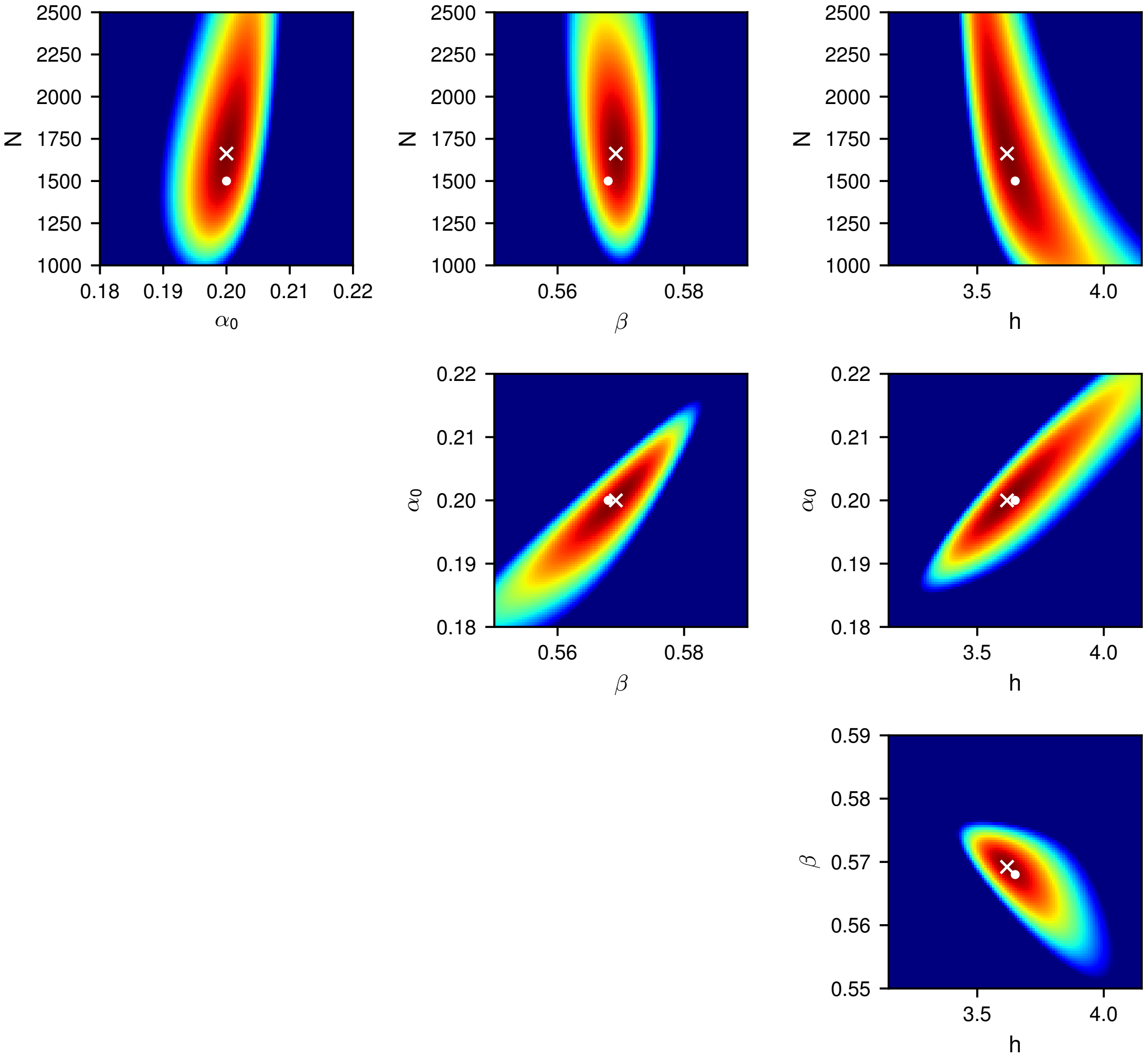


**Figure S9**: **Likelihood distribution for the SRG model with $h = 3.65$.** Likelihood distribution for inference of all pairs of parameters for the SRG model, using the optimized prefactors. For each plot, all other parameters are fixed to their MLE. The MLE is marked with a white $\times$ and the true parameter values are marked with a white $\bullet$. Colors show $\log_{10}[\mathrm{L}]$ and range from $-1.0{\times}10^5$ (blue) to $0$ (red).

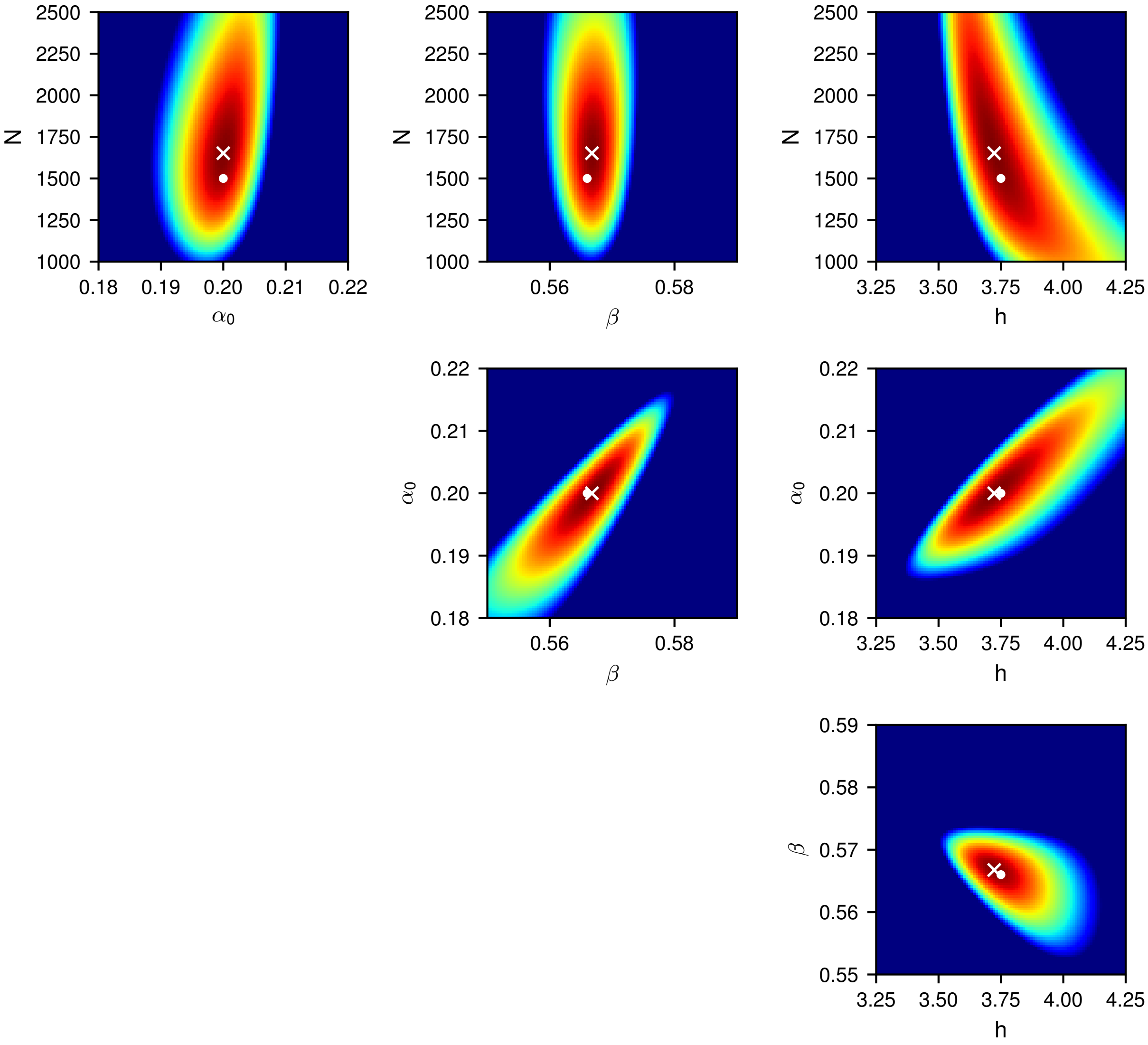


**Figure S10**: **Likelihood distribution for the SRG model with $h = 3.75$.** Likelihood distribution for inference of all pairs of parameters for the SRG model, using the optimized prefactors. For each plot, all other parameters are fixed to their MLE. The MLE is marked with a white $\times$ and the true parameter values are marked with a white •. Colors show $\log_{10}[\mathrm{L}]$ and range from $-1.0\times10^5$ (blue) to $0$ (red).

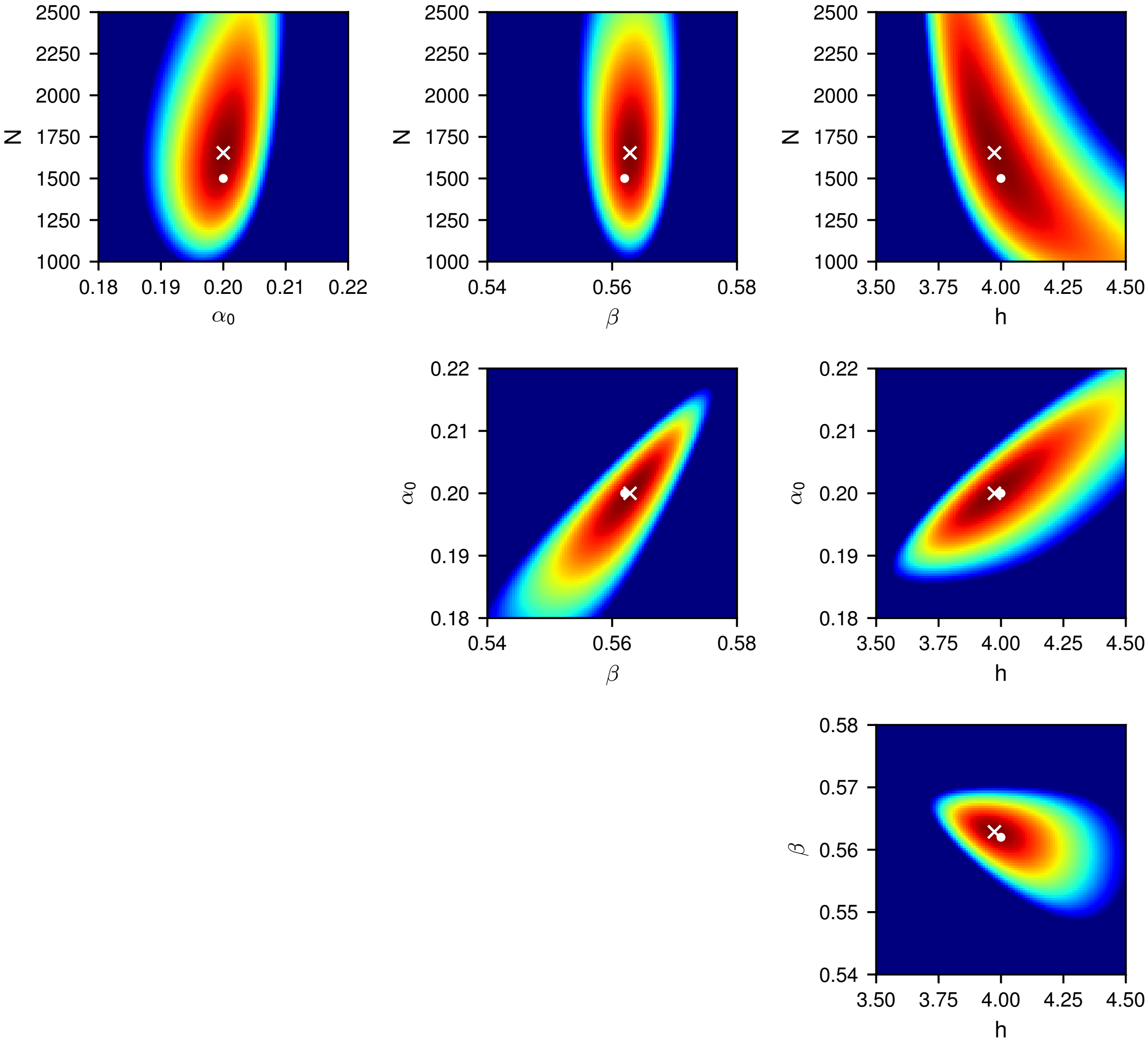


**Figure S11**: **Likelihood distribution for the SRG model with $h = 4.0$.** Likelihood distribution for inference of all pairs of parameters for the SRG model, using the optimized prefactors. For each plot, all other parameters are fixed to their MLE. The MLE is marked with a white $\times$ and the true parameter values are marked with a white $\bullet$. Colors show $\log_{10}[\mathrm{L}]$ and range from $-1.0\times10^5$ (blue) to $0$ (red).

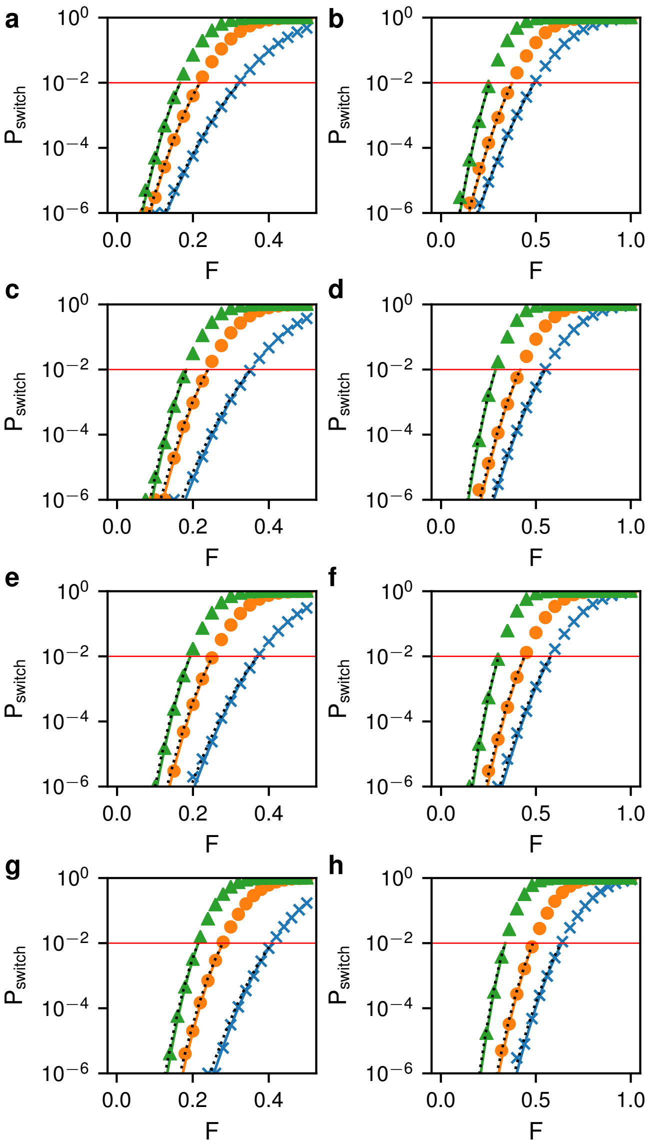


**Figure S12**: **Perturbation effect on the SRG model.** (a) Change in switching probability vs perturbation strength $F$ for the *low* to *high* switch with $h = 3.5$. Shown are the numerical solution (symbols), theory with MLE parameters (solid lines), and theory with true parameters and 0.15 prefactor (dotted lines). Colors give the perturbation time $T$: 0.5 (blue $\times$), 0.75 (orange $\circ$), 1.0 (green $\triangle$). (b) Change in switching probability vs perturbation strength for the *high* to *low* switch with perturbation times 0.75 (blue $\times$), 1.0 (orange $\circ$), and 1.5 (green $\triangle$). Here the prefactor for the true parameter line was 0.2. (c+d) As in (a+b) except for $h = 3.65$. (e+f) $h = 3.75$. (g+h) $h = 4.0$.

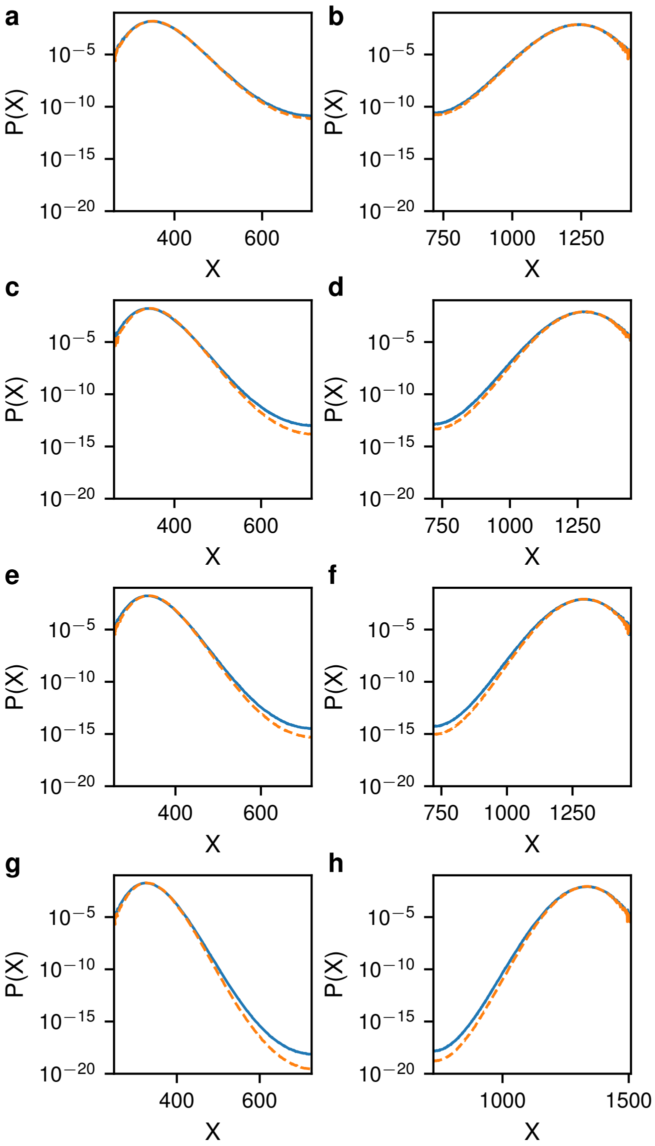


**Figure S13**: **Comparison of actual and inferred probability distributions for the self-regulating gene.** (a+b) The actual (solid blue) and inferred (dashed orange) PDFs for the *low* state (a) and *high* state (b) for $h = 3.5$. (c+d) As in (a+b) except for $h = 3.65$. (e+f) $h = 3.75$. (g+h) $h = 4.0$.

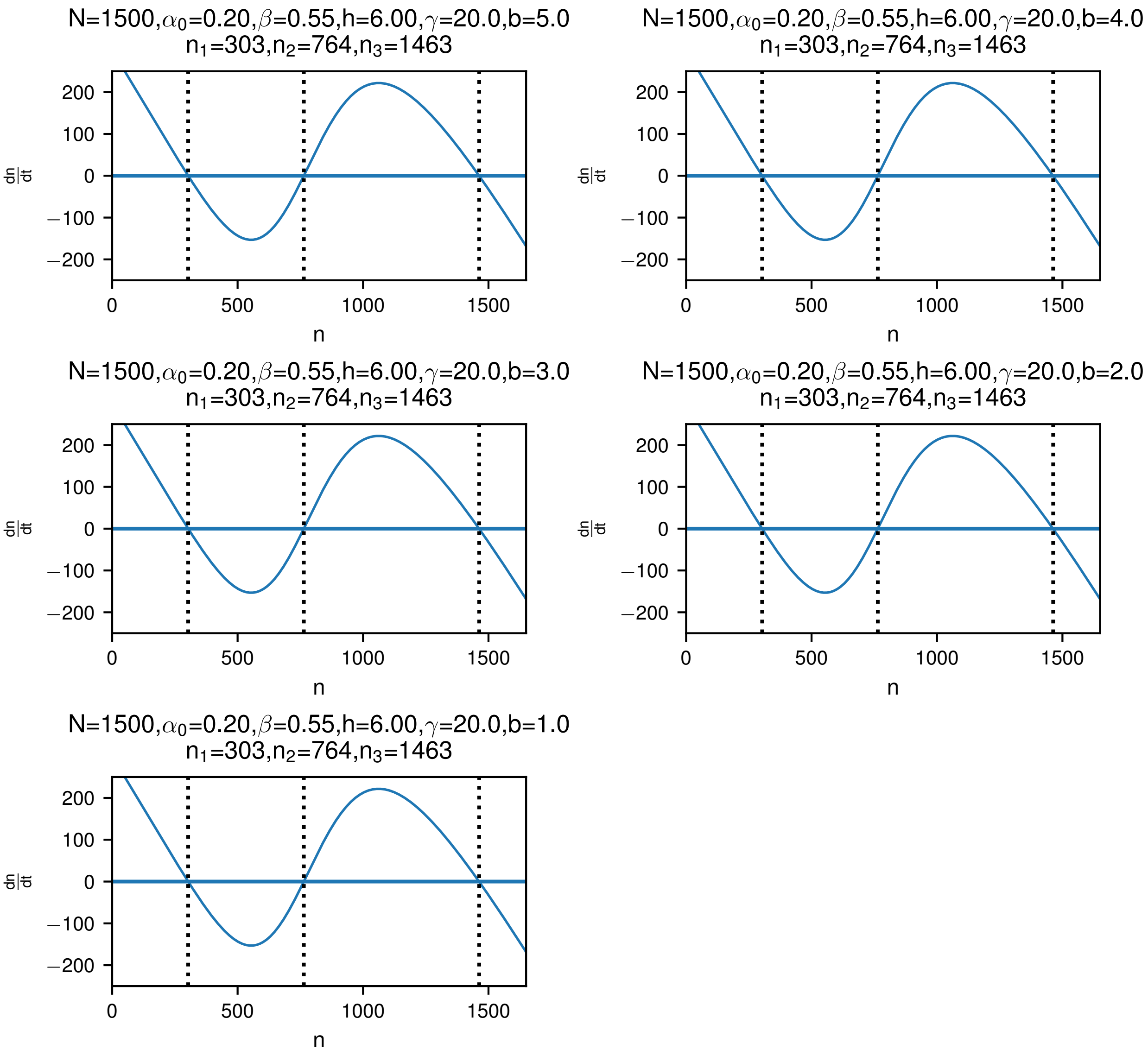


**Figure S14**: **Deterministic rate equations for the mrna-protein model.** Value of $\frac{dn}{dt}$ as a function of $n$ for the deterministic model of the mrna-protein switch. The positions of the three fixed points are given by the dotted lines. Parameters for each panel are as indicated.

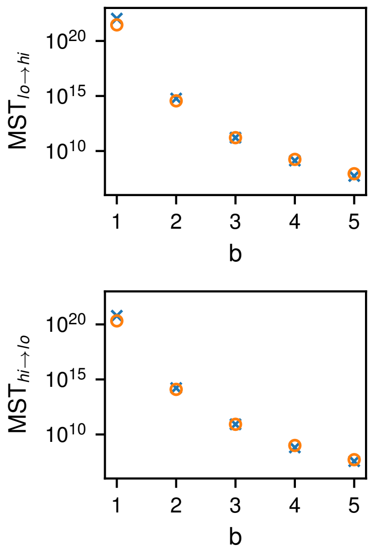


**Figure S15**: **Mean switching times for the mrna-protein model.** (top) The MST to go from the $low$ state to the $high$ state vs $b$ calculated from numerical simulations (blue $\times$) and WKB theory (orange $\circ$) as given by Eq. (24) in the main text. The WKB points are multiplied by a constant preexponential factor of 37.33. (bottom) The same for the $high$ to $low$ state with a preexponent of 14.11. All other parameters are as in Figure S14.

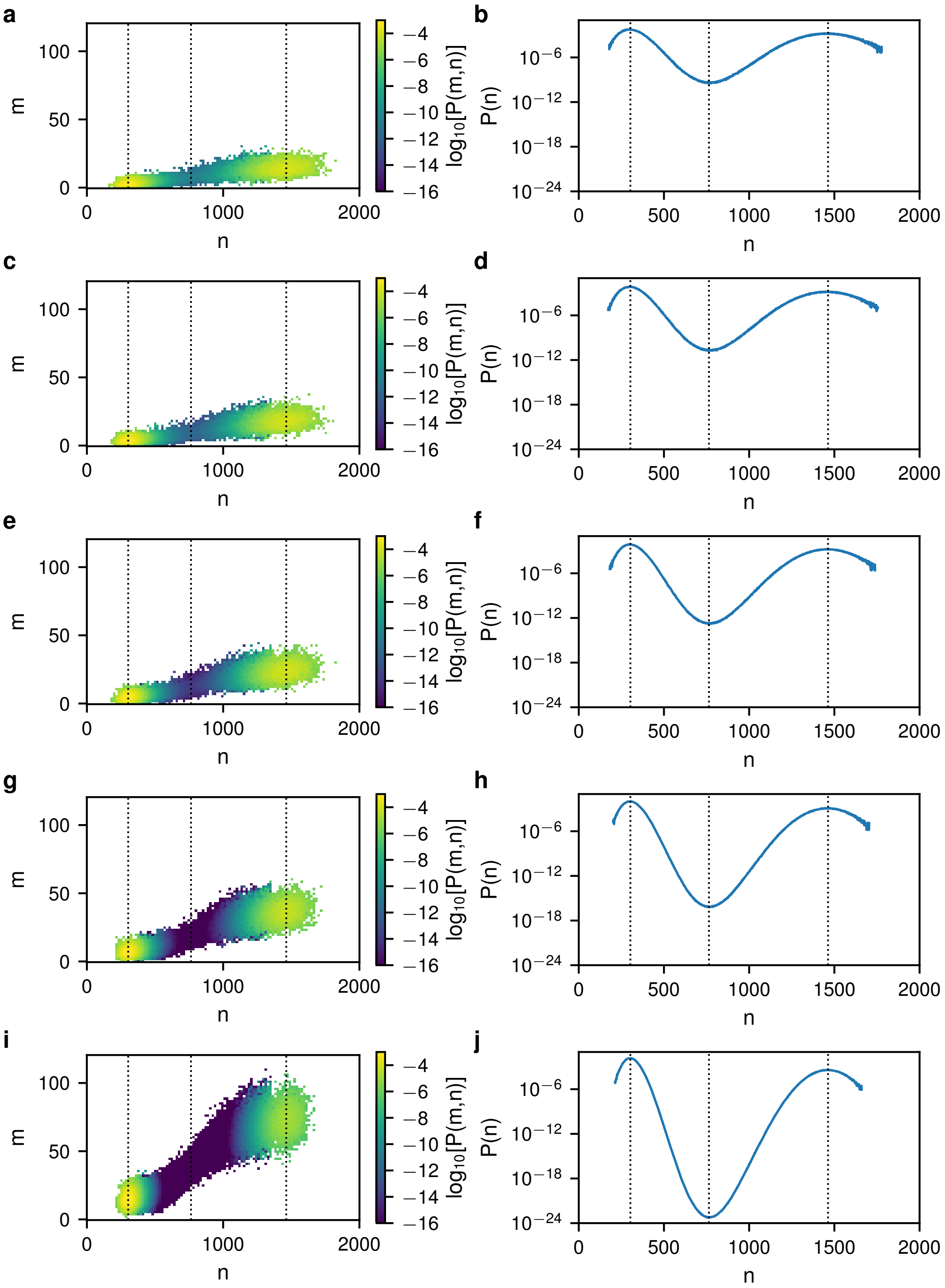


**Figure S16**: **Stationary probability distributions for the mrna-protein model.** (a) The joint probability density for a given number of mrna (m) and protein (n) molecules for the mrna-protein switch with $b = 5.0$. (b) The marginal probability density for only the protein count (n) with $b = 5$. (c+d) As in (a+b) except for $4 = 2$. (e+f) $b = 3$. (g+h) $b = 2$. (i+j) $b = 1$.

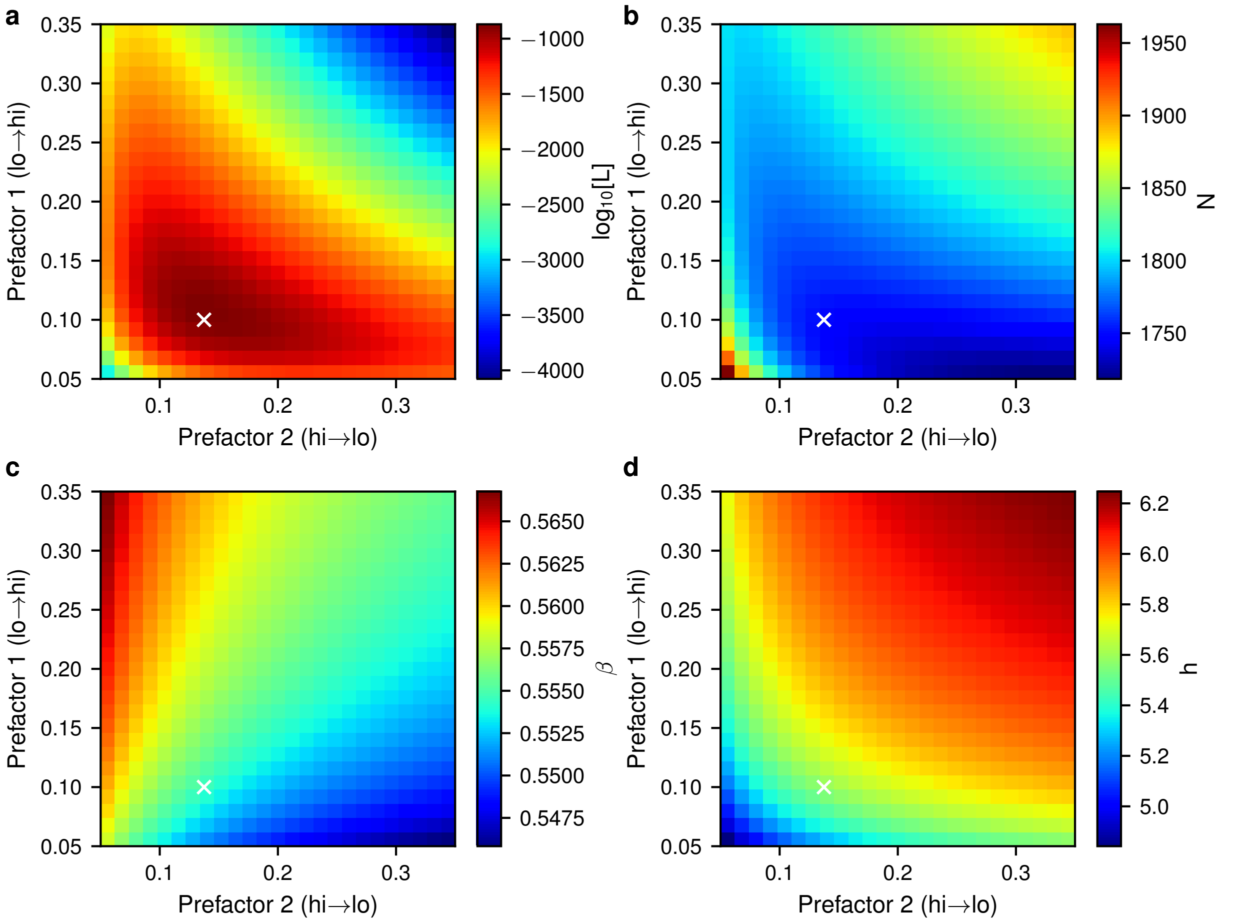


**Figure S17**: **Prefactor dependence during maximum likelihood fitting of the mrna-protein model with $b = 5$.** (a) The maximum likelihood score obtained for each prefactor pair during fitting. Fitting was performed with $\alpha$ and $b$ fixed to their true value, as described in the main text, using simulation data obtained from the mrna-protein model with parameters $N = 1500$, $\alpha_0 = 0.2$, $\beta = 0.55$, $h = 6.0$, $\gamma = 20$, $b = 5$. The prefactor pair with the highest likelihood score is marked with a white $\times$ in each panel. (b-d) The dependence of the maximum likelihood estimate for the parameters $N$, $\beta$, and $h$, respectively, on the prefactors.

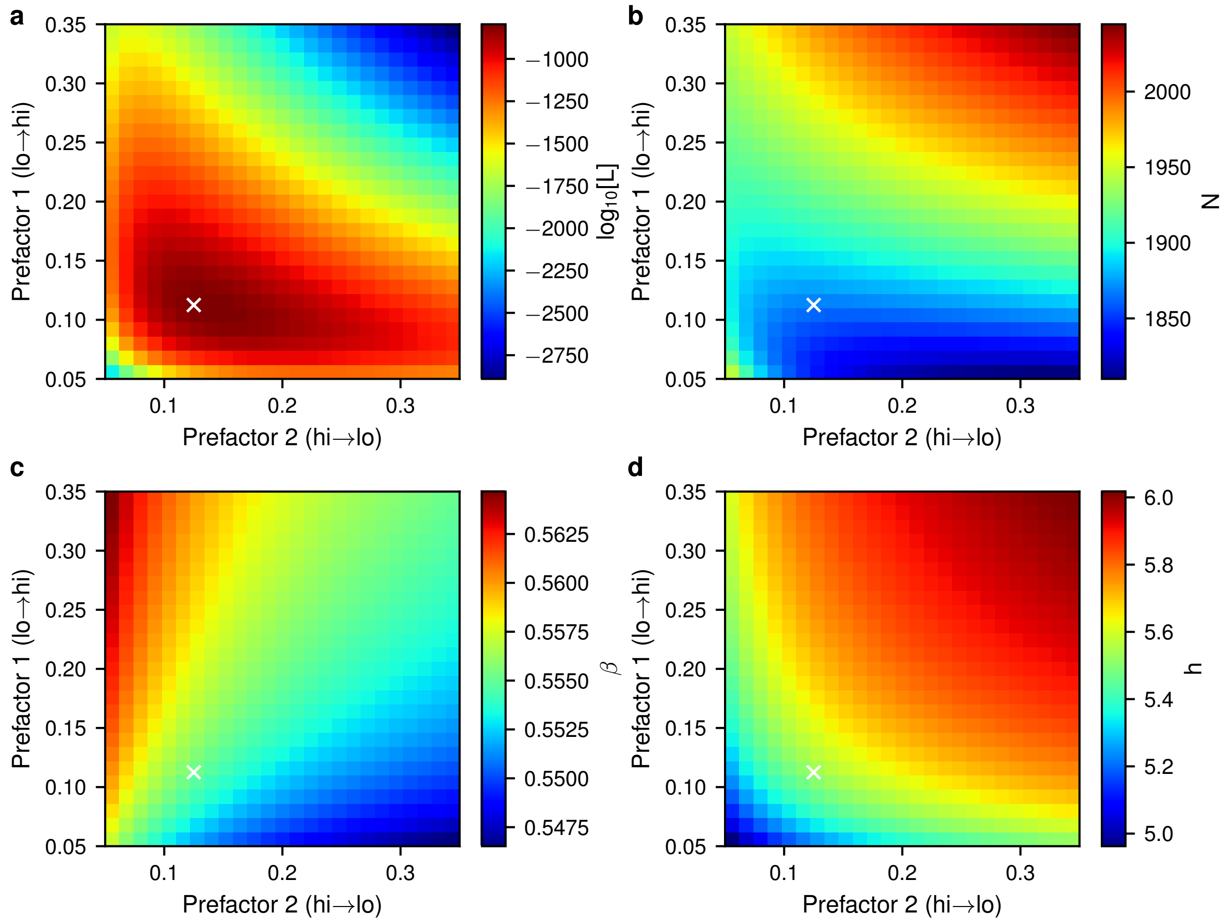


**Figure S18**: **Prefactor dependence during maximum likelihood fitting of the mrna-protein model with $b = 4$.** (a) The maximum likelihood score obtained for each prefactor pair during fitting. Fitting was performed with $\alpha$ and $b$ fixed to their true value, as described in the main text, using simulation data obtained from the mrna-protein model with parameters $N = 1500$, $\alpha_0 = 0.2$, $\beta = 0.55$, $h = 6.0$, $\gamma = 20$, $b = 4$. The prefactor pair with the highest likelihood score is marked with a white $\times$ in each panel. (b-d) The dependence of the maximum likelihood estimate for the parameters $N$, $\beta$, and $h$, respectively, on the prefactors.

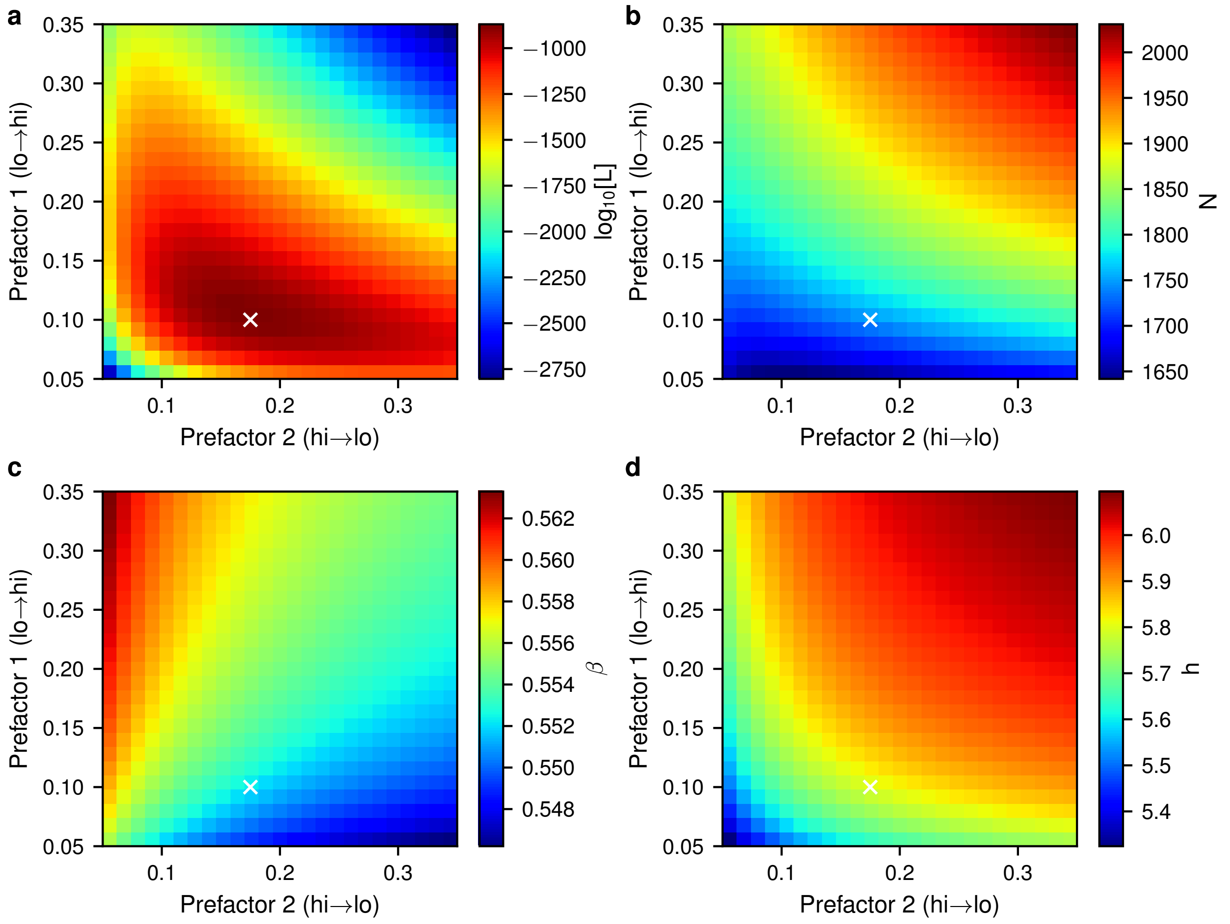


**Figure S19**: **Prefactor dependence during maximum likelihood fitting of the mrna-protein model with $b = 3$.** (a) The maximum likelihood score obtained for each prefactor pair during fitting. Fitting was performed with $\alpha$ and $b$ fixed to their true value, as described in the main text, using simulation data obtained from the mrna-protein model with parameters $N = 1500$, $\alpha_0 = 0.2$, $\beta = 0.55$, $h = 6.0$, $\gamma = 20$, $b = 3$. The prefactor pair with the highest likelihood score is marked with a white $\times$ in each panel. (b-d) The dependence of the maximum likelihood estimate for the parameters $N$, $\beta$, and $h$, respectively, on the prefactors.

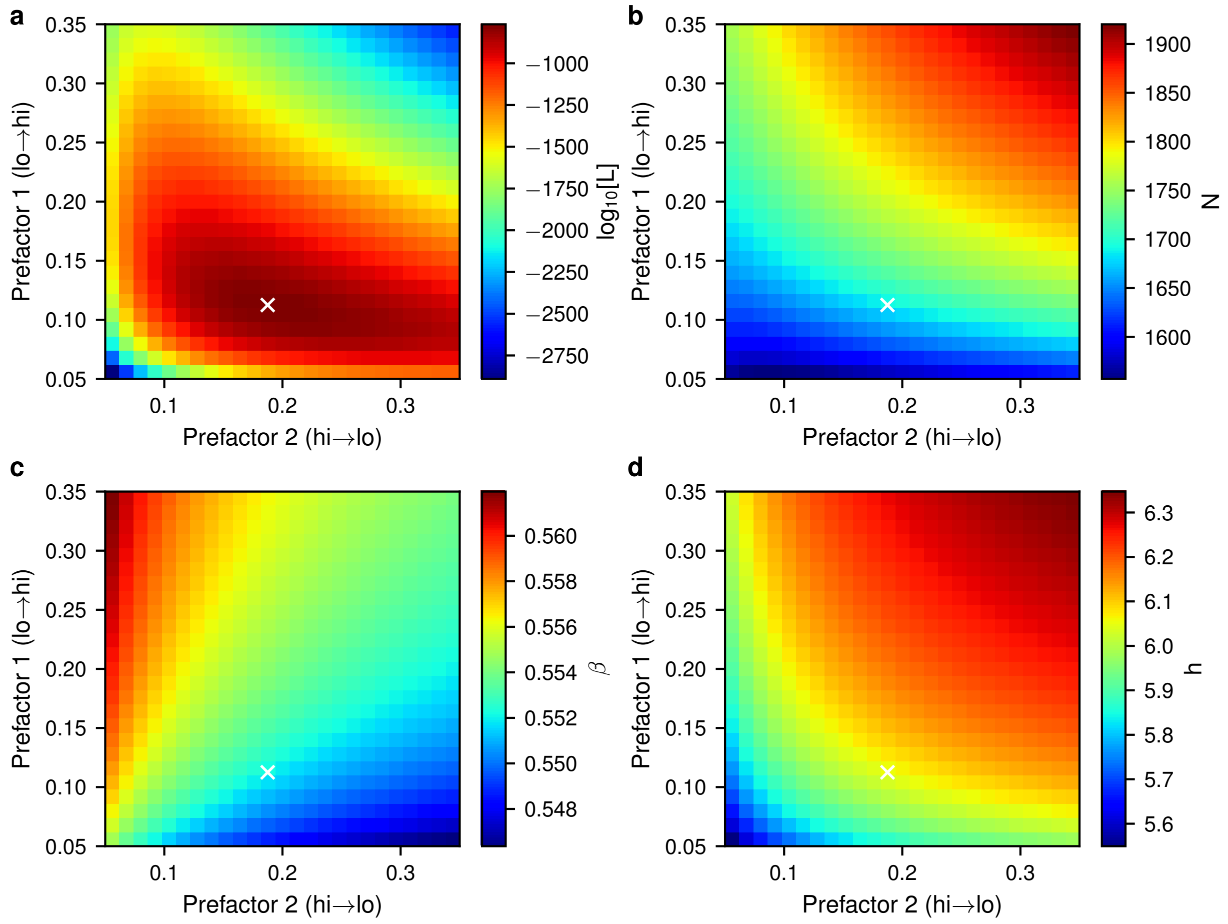


**Figure S20**: **Prefactor dependence during maximum likelihood fitting of the mrna-protein model with $b = 2$.** (a) The maximum likelihood score obtained for each prefactor pair during fitting. Fitting was performed with $\alpha$ and $b$ fixed to their true value, as described in the main text, using simulation data obtained from the mrna-protein model with parameters $N = 1500$, $\alpha_0 = 0.2$, $\beta = 0.55$, $h = 6.0$, $\gamma = 20$, $b = 2$. The prefactor pair with the highest likelihood score is marked with a white $\times$ in each panel. (b-d) The dependence of the maximum likelihood estimate for the parameters $N$, $\beta$, and $h$, respectively, on the prefactors.

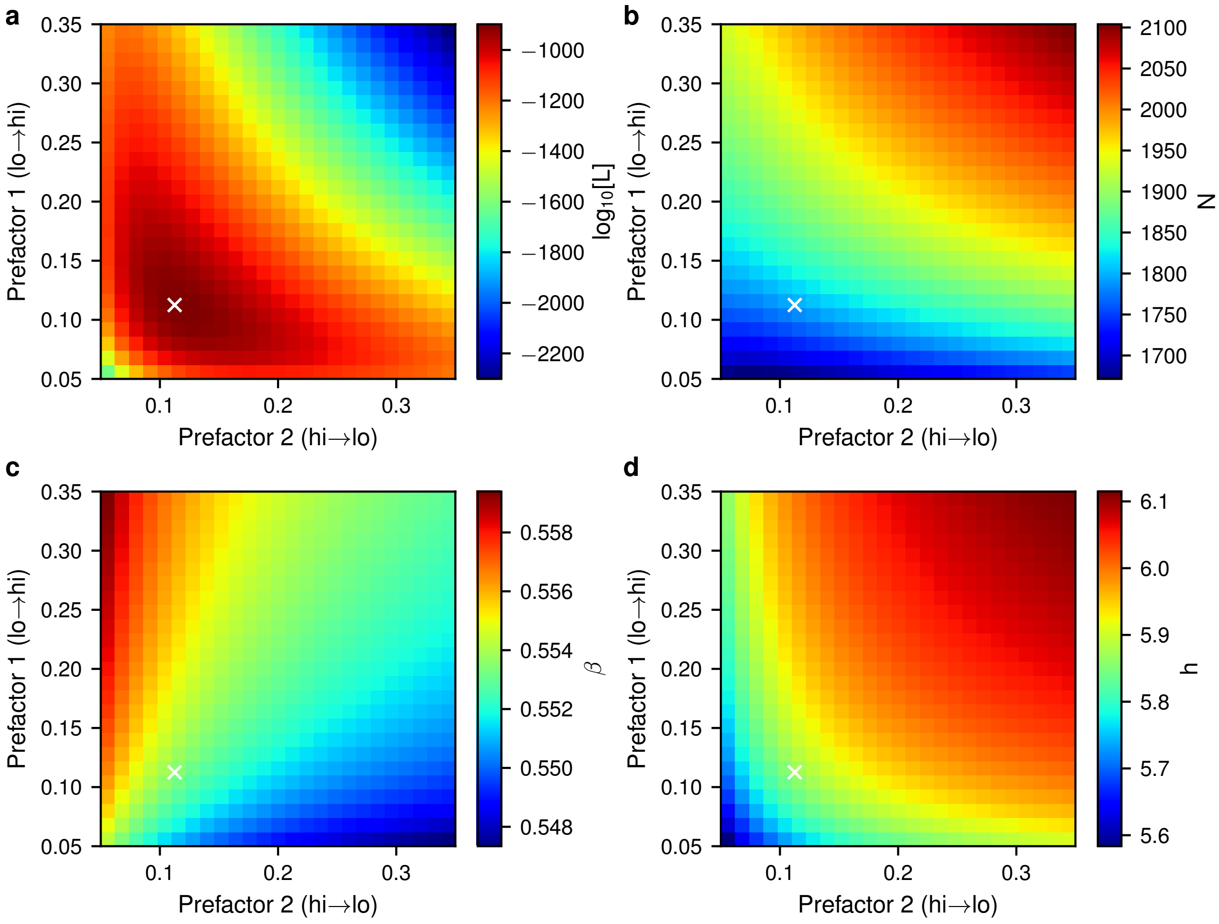


**Figure S21**: **Prefactor dependence during maximum likelihood fitting of the mrna-protein model with** $b = 1$**.** (a) The maximum likelihood score obtained for each prefactor pair during fitting. Fitting was performed with $\alpha$ and $b$ fixed to their true value, as described in the main text, using simulation data obtained from the mrna-protein model with parameters $N = 1500$, $\alpha_0 = 0.2$, $\beta = 0.55$, $h = 6.0$, $\gamma = 20$, $b = 1$. The prefactor pair with the highest likelihood score is marked with a white $\times$ in each panel. (b-d) The dependence of the maximum likelihood estimate for the parameters $N$, $\beta$, and $h$, respectively, on the prefactors.

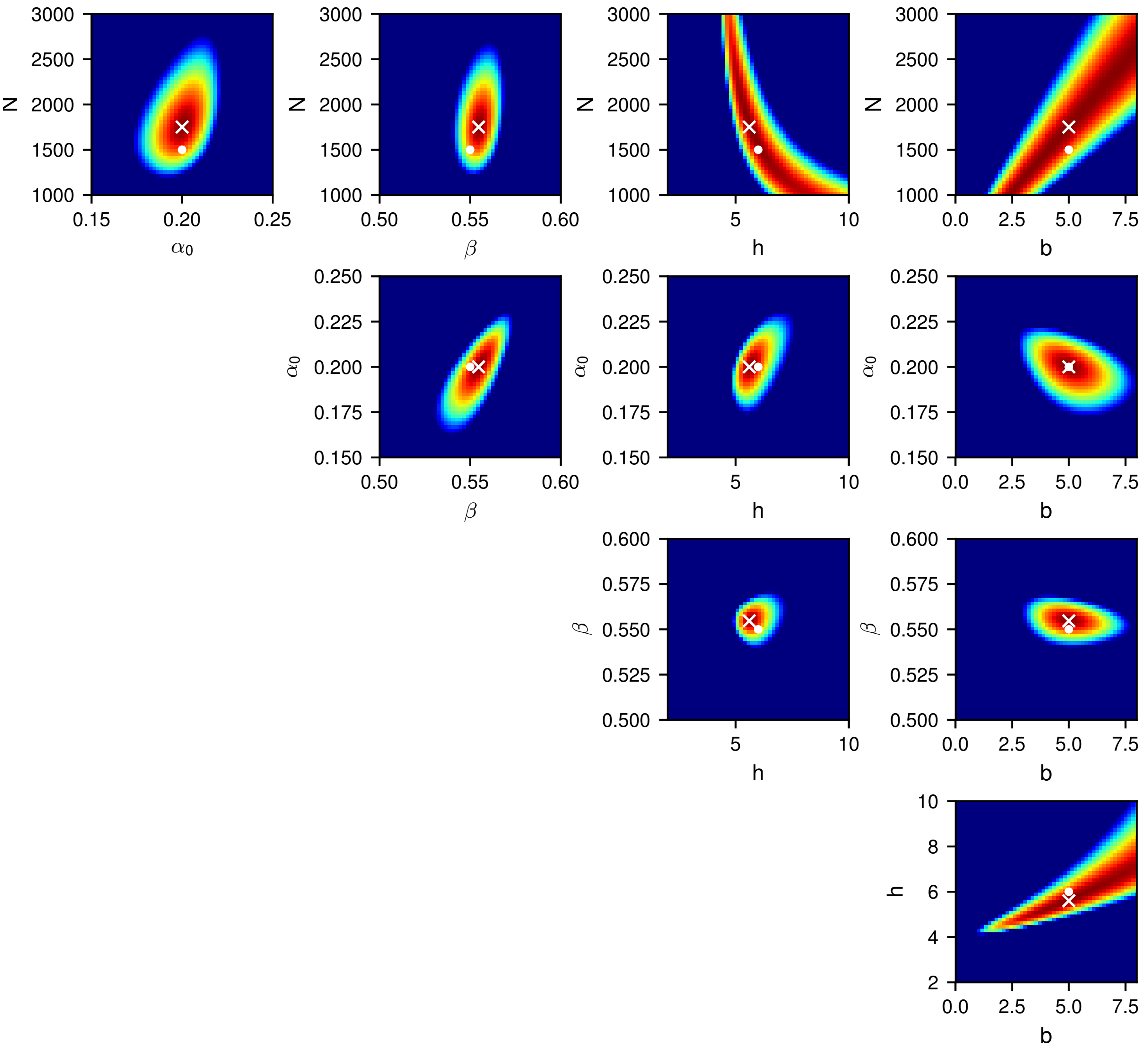


**Figure S22**: **Likelihood distribution for the mrna-protein model with $b = 5$.** Likelihood distribution for inference of all pairs of parameters for the mrna-protein model, using the optimized prefactors. For each plot, all other parameters are fixed to their MLE. The MLE is marked with a white $\times$ and the true parameter values are marked with a white •. Colors show $\log_{10}[\mathrm{L}]$ and range from $-1.0{\times}10^5$ (blue) to $0$ (red).

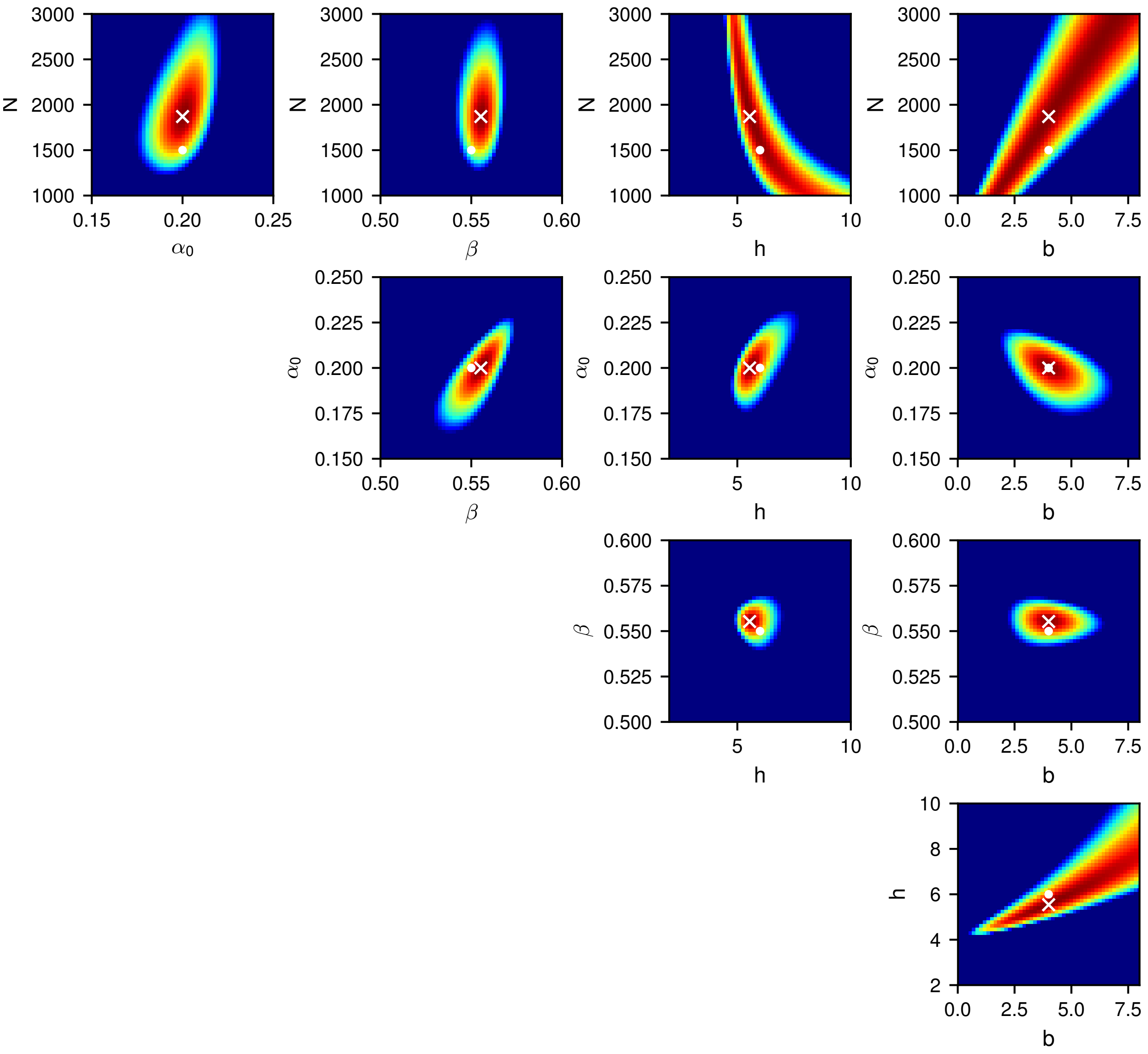


**Figure S23**: **Likelihood distribution for the mrna-protein model with $b = 4$.** Likelihood distribution for inference of all pairs of parameters for the mrna-protein model, using the optimized prefactors. For each plot, all other parameters are fixed to their MLE. The MLE is marked with a white $\times$ and the true parameter values are marked with a white $\bullet$. Colors show $\log_{10}[\mathrm{L}]$ and range from $-1.0\times10^5$ (blue) to $0$ (red).

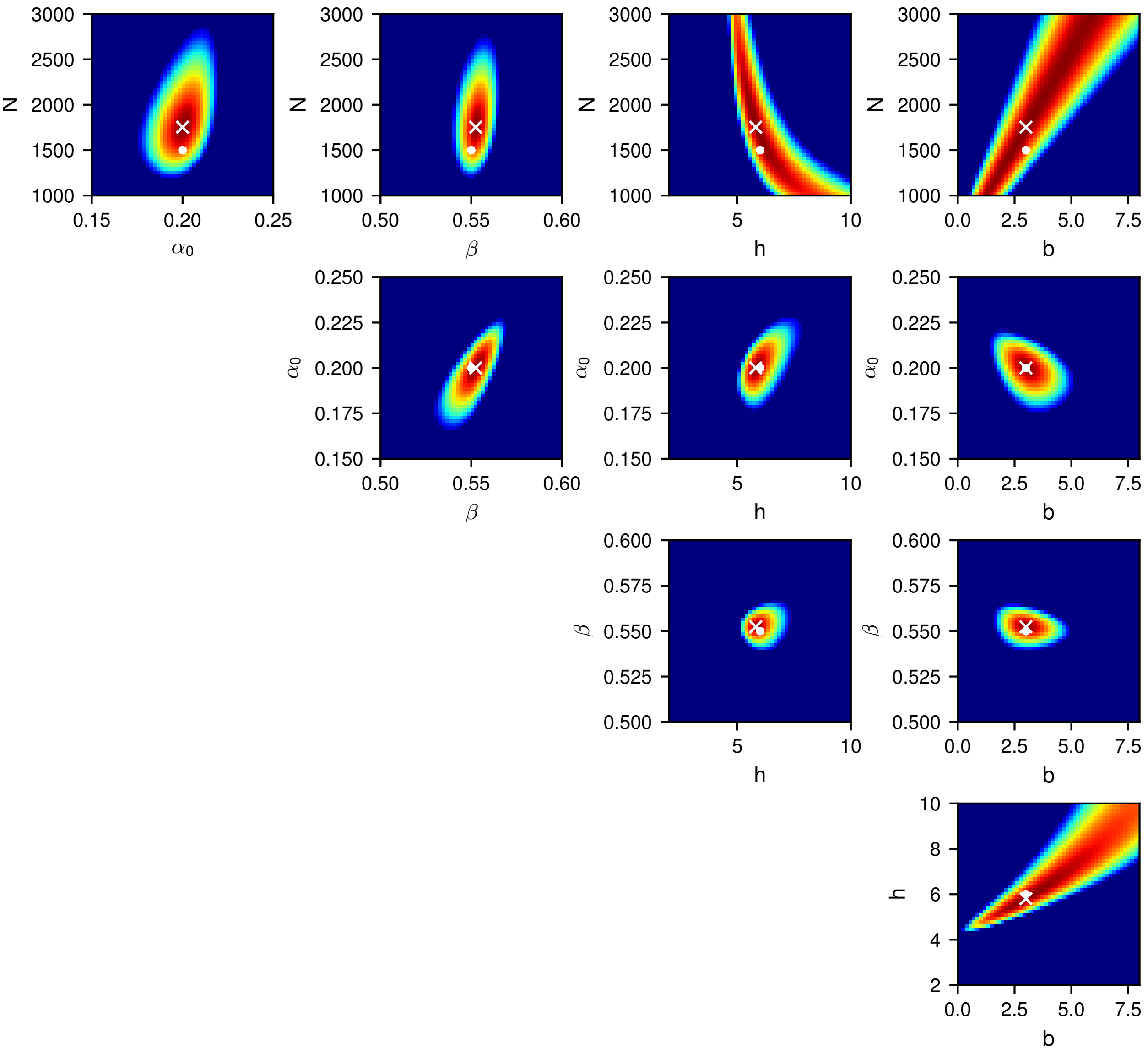


**Figure S24**: **Likelihood distribution for the mrna-protein model with $b = 3$.** Likelihood distribution for inference of all pairs of parameters for the mrna-protein model, using the optimized prefactors. For each plot, all other parameters are fixed to their MLE. The MLE is marked with a white $\times$ and the true parameter values are marked with a white •. Colors show $\log_{10}[\mathrm{L}]$ and range from $-1.0\times10^5$ (blue) to $0$ (red).

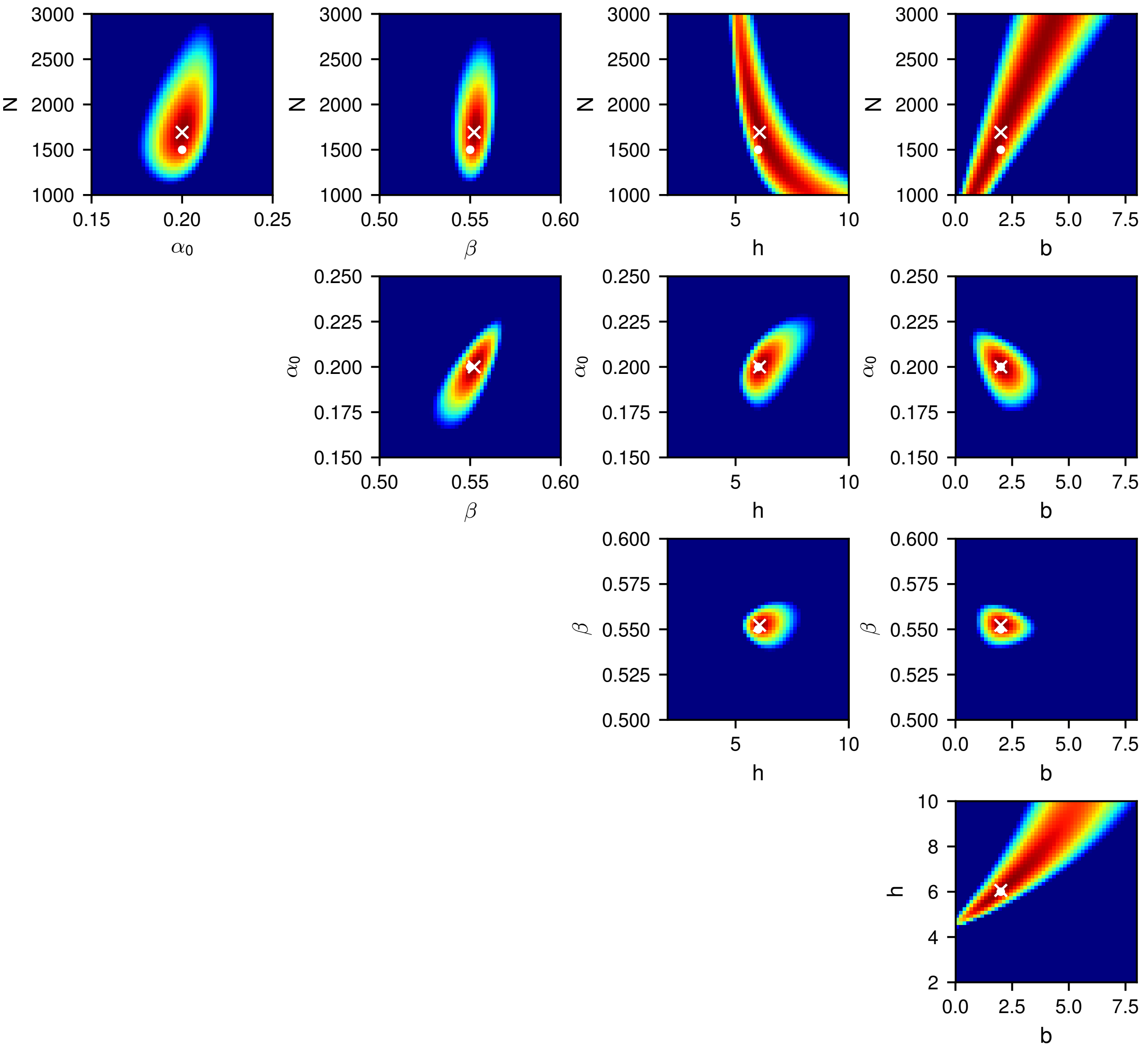


**Figure S25**: **Likelihood distribution for the mrna-protein model with $b = 2$.** Likelihood distribution for inference of all pairs of parameters for the mrna-protein model, using the optimized prefactors. For each plot, all other parameters are fixed to their MLE. The MLE is marked with a white $\times$ and the true parameter values are marked with a white •. Colors show $\log_{10}[\mathrm{L}]$ and range from $-1.0{\times}10^5$ (blue) to $0$ (red).

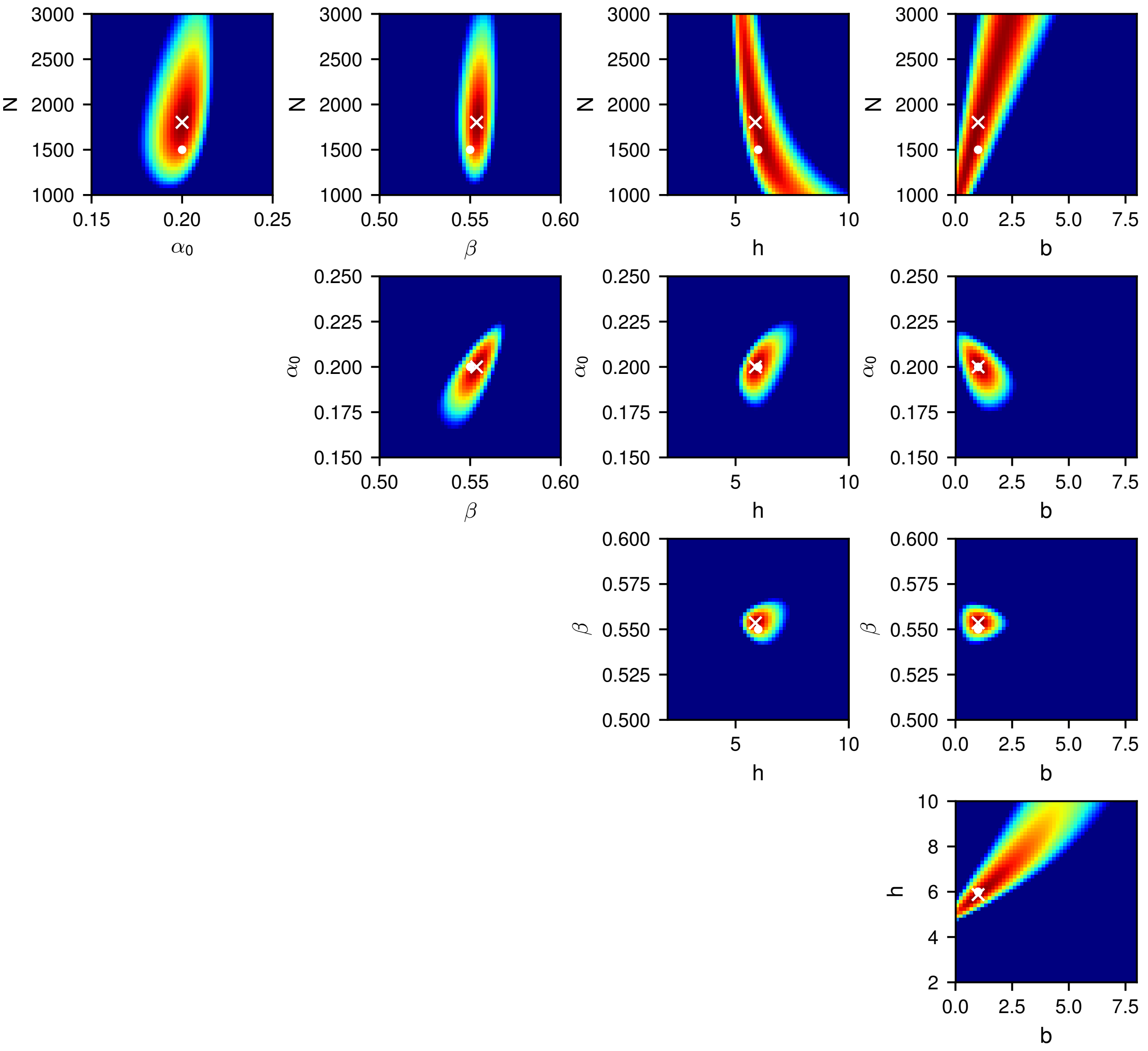

**Figure S26**: **Likelihood distribution for the mrna-protein model with $b = 1$.** Likelihood distribution for inference of all pairs of parameters for the mrna-protein model, using the optimized prefactors. For each plot, all other parameters are fixed to their MLE. The MLE is marked with a white $\times$ and the true parameter values are marked with a white •. Colors show $\log_{10}[\mathrm{L}]$ and range from $-1.0{\times}10^5$ (blue) to $0$ (red).

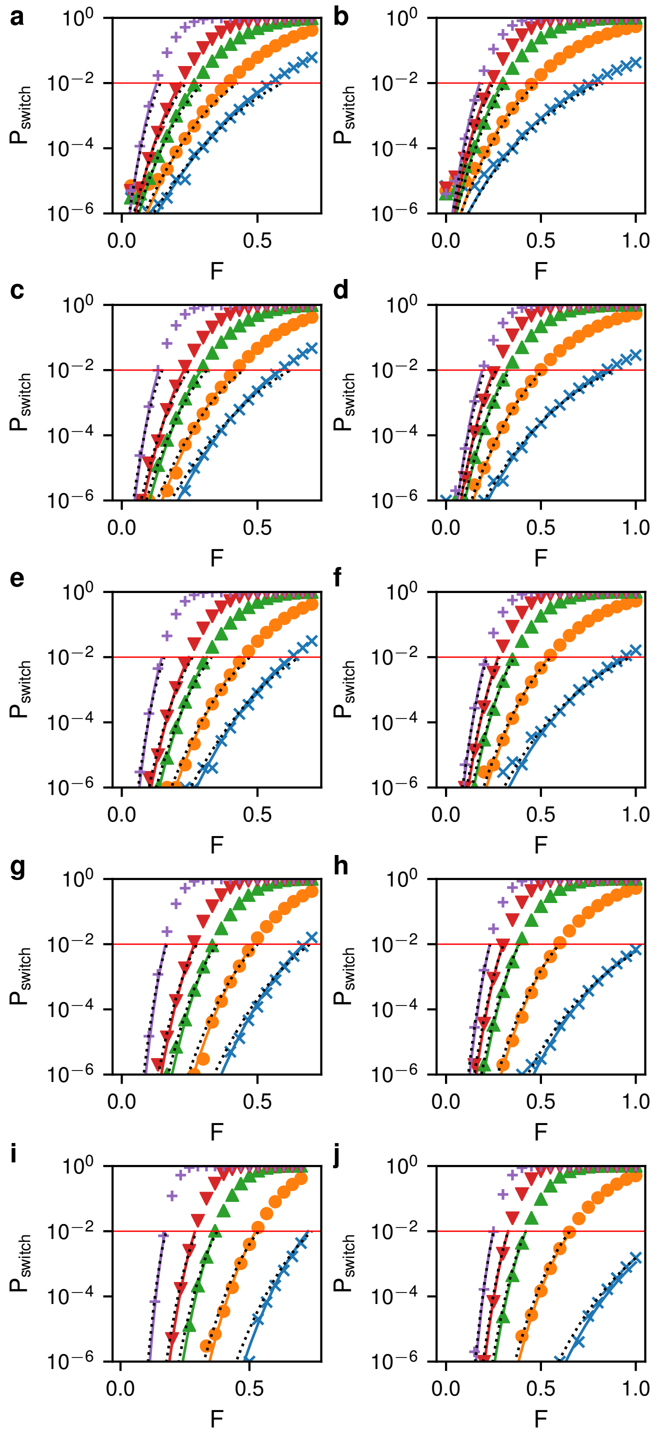


**Figure S27**: **Perturbation effect on the mrna-protein model.** (a) Change in switching probability vs perturbation strength $F$ for the *low* to *high* switch with $b = 5$. Shown are the numerical solution (symbols), theory with MLE parameters (solid lines), and theory with true parameters and 0.05 prefactor (dotted lines). Colors give the perturbation time $T$: 0.35 (blue $\times$), 0.5 (orange $\circ$), 0.75 (green $\triangle$), 1.0 (red $\bigtriangledown$), 2.0 (purple $+$). (b) Change in switching probability vs perturbation strength for the *high* to *low* switch with perturbation times 1.0 (blue $\times$), 1.5 (orange $\circ$), 2.25 (green $\triangle$), 3.0 (red $\bigtriangledown$), 4.5 (purple $+$). Here the prefactor for the true parameter line was 0.15. (c+d) As in (a+b) except for $b = 4$. (e+f) $b = 3$. (g+h) $b = 2$ (i+j) $b = 1$.

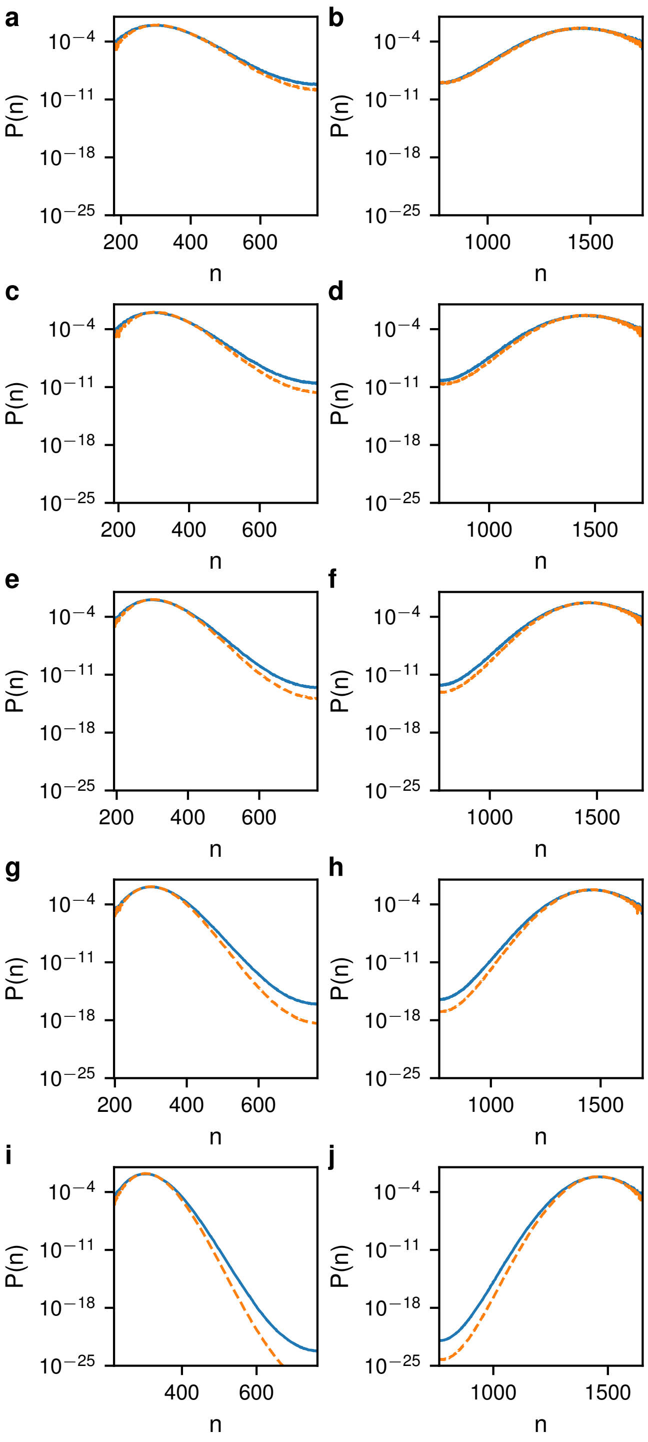


**Figure S28**: **Comparison of actual and inferred probability distributions for the mrna-protein model.** (a+b) The actual (solid blue) and inferred (dashed orange) PDFs for the *low* state (a) and *high* state (b) for $b = 5$. (c+d) As in (a+b) except for $b = 4$. (e+f) $b = 3$. (g+h) $b = 2$ (i+j) $b = 1$.

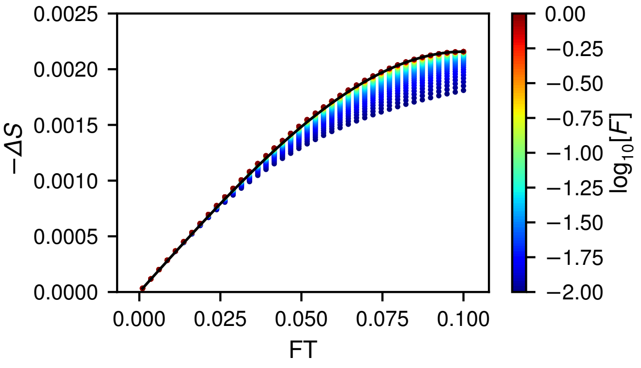


**Figure S29**: **Dependence of change in the switching barrier on impulse near bifurcation.** Distribution of the perturbed switching barrier $\Delta S = \mathcal{S}^{lh} - \mathcal{S}_0^{lh}$ versus the total applied impulse $FT$ for the SRG model. Each symbol represents a perturbation with a different $F$, which is given by the color. The solid line shows the bifurcation theory given in Eq. (44) in the main text. The parameters were $\alpha_0 = 0.2$, $\beta = 0.501$ and $h = 4$
